# Disparate introduction histories but similar climatic distribution patterns among congeneric invasive anurans

**DOI:** 10.64898/2026.06.08.730926

**Authors:** Andrew J. Mularo, Jong Yoon Jeon, Jackson T. Kirkwood, Ximena E. Bernal

**Affiliations:** Department of Biological Sciences, Purdue University, West Lafayette, Indiana, USA; Department of Forestry and Natural Resources, Purdue University, West Lafayette, Indiana, USA; Smithsonian Tropical Research Institute, Apartado, Republic of Panama

**Keywords:** Climate change, ecological niche modeling, *Eleutherodactylus*, invasive species, range expansion

## Abstract

Commonly shared patterns of introduction and spread into new environmental conditions are often poorly understood, even though a better understanding of invasion history and niche dynamics among closely related invasive species could give practitioners valuable information to prevent and mitigate the impact of biological invasions. For this study, we investigate the invasion history and niche patterns among congeneric invasive species. We synthesize public occurrence data for five invasive alien anurans (*Eleutherodactylus coqui, E. planirostris, E. johnstonei, E. antillensis, and E. martinicensis*) to reconstruct their historic introductions and evaluate evidence for climatic niche shifts between their native and established non-native ranges. By pairing these data with current and future climate projections, we compare patterns of range shifts under future climate scenarios. Our results highlight different temporal and geographic introduction histories in invasive *Eleutherodactylus*, but a strong signal of colonizing broader invasive climatic niches, specifically into colder environmental conditions. Under future climate scenarios, suitable habitats for most of the non-native regions are likely to increase, although this increase is restricted under scenarios with high greenhouse gas emissions. Our results reveal that despite different invasion histories, the ability to spread into colder regions may be a conserved trait among the most widespread *Eleutherodactylus* anurans. This study ultimately shows that commonalities among closely related invasive species can provide clues about their ability to expand into areas with particular abiotic conditions, a pattern likely to be widespread, offering a potentially valuable opportunity to deploy targeted prevention strategies.

## Introduction

Developing the ability to accurately predict invasion potential is one of the most pressing issues in the field of invasion biology. Often, the risk of invasion is determined by reconstructing the environmental niche of species to determine the likelihood of species occurrence based on the environmental conditions in a region. For many biological invasions, environmental niches are considered largely conserved between a species’ native and invasive range (Townsend Peterson 2011; Strubbe et al. 2013; Liu et al. 2017). At the same time, however, there are many instances of climatic niche expansion when a species colonizes a new range (Tingley et al. 2014; Rödder et al. 2017). This shift may occur, for example, when species have significant biotic or geographic limits to dispersal in their native range but are released from those constraints when introduced into a new environment (Alexander and Edwards 2010). Niche expansion is also more likely the longer an invasive species spends in a novel environment, as more time allows for recovery of lost genetic variation, facilitating adaptation or phenotypic plasticity towards novel conditions (Alexander and Edwards 2010). As such, island endemics with a long invasion history frequently show niche expansions when introduced into a new range (Li et al. 2014). Given their unpredictability, niche expansions complicate efforts to accurately anticipate the likelihood that species will successfully invade new environments, and invasion patterns that can be generalized among species in “high-risk” taxa remain rare. There remains a crucial need to assess the complexities of invasion history and niche dynamics among closely related invasive species to ultimately discover shared, predictable patterns of invasion.

The reconstruction of a species’ invasion history and niche dynamics often require consulting a variety of sources (i.e. museum specimens, published literature, online databases) (Suarez et al. 2001; Saul et al. 2017; Hiller and Haelewaters 2019; Tedeschi et al. 2022) that ultimately allow estimations of species distribution (Galera and Sudnik-Wójcikowska 2010; Kannan et al. 2013; Dyer et al. 2017, Biancolini et al. 2021), identification of common geographic regions of introduction and mechanisms of spread (Lehan et al. 2013; Zenni 2013; Mularo et al. 2023), as well as projections of future introductions (Ficetola et al. 2007; Bellard et al. 2013; Li et al. 2016; Biancolini et al. 2024). Museum collections, in particular, are commonly used to study the historical trajectory of biological invasions (Pysek and Prach 1993; Suarez et al. 2001; Delisle et al. 2003; Suarez and Tsutsui 2004; Kannan et al. 2013). Additionally, the use of community science through platforms such as iNaturalist (http://www.inaturalist.org) has become increasingly popular for supplementing museum records, allowing real-time and large-scale observations of introduced species (e.g. Hiller and Haelewaters; 2019) unavailable in museums or other research data. Perhaps most importantly, geo-referenced records can be correlated with other forms of environmental data to estimate potential species distributions (Newbold 2010), compare the environmental conditions between native and non-native ranges (Liu et al. 2020; Bates and Bertelsmeier 2021), and generate projections of the potential future range of invasive species (Rödder and Weinsheimer 2009; Sales et al. 2021).

Despite the importance of understanding the historical context of invasions, poorly documented invasion histories are common among many organisms, an issue that can lead to misidentifications of future areas of potential spread. In the anuran genus *Eleutherodactylus*, five island endemic species – *Eleutherodactylus antillensis, E, coqui*, *E. johnstonei, E. martinicensis,* and *E. planirostris* – native to the Caribbean have been transported, likely through the nursery plant trade (Christy et al. 2007a,b; Kraus 2009; Powell et al. 2011), to many regions across the globe. The history of invasion of all five species, however, remains poorly documented, as early records rely on anecdotal accounts with considerable disagreement on the establishment status in several non-native countries (Lever 2003; Kraus 2009; Powell 2011). *Eleutherodactylus* are small frogs (<35mm) and undergo direct development (Townsend and Stewart 1985), laying their eggs on land in humid microhabitats (e.g. large fallen leaves, plant pots; Westrick et al. 2022) - features that reduce detectability and facilitate transport and establishment in new regions. These cryptic introductions are concerning, as members of this genus can pose serious ecological and economic risks in non-native areas.

*Eleutherodactylus coqui* is a well-recognized invader in Hawai’i and, due to its loud calls, its presence can reduce property values (Beard et al. 2009). Indeed, *E. coqui* is considered among the 100 worst invasive alien species in the world (Simberloff and Rejmanek 2019) and is categorized as being of “moderate concern” by the IUCN in the Environmental Impact Classification for Alien Taxa (EICAT) (Global Invasive Species Database 2025). The impacts of the other four species are less known, although it has been proposed that some compete for resources with various native anuran species in their introduced range (Kaiser and Henderson 1994; Meshaka *et al*. 2004). While *E. johnstonei* is considered of ‘minimal concern’, *E. antillensis* and *E. martinicensis* are considered ‘data deficient’, and *E. planirostris* has yet to be assigned to a category by the EICAT (Global Invasive Species Database 2025). Limited knowledge about these species highlights their poorly understood potential spread and impact. Besides their direct impacts, the introduction of anuran species raises concern due to the potential of spreading the chytrid fungus *Batrachochytrium dendrobatidis* (*Bd*), which is one of the major causes of the global amphibian decline (Fisher and Garner 2020). These anuran species can also facilitate the occurrence of other pathogens, such as antibiotic-resistant microbes (Abreu et al. 2025). Previous studies have projected the potential spread of *E. coqui*, *E. planirostris*, and *E. johnstonei* under current and future climate conditions (Rödder 2009a,b; Mu et al. 2022), providing a valuable opportunity to refine the historical context and examine common ecological patterns that may promote the successful establishment of species from this genus in non-native regions.

For this study, we compare the invasion history and niche breadth among the invasive alien *Eleutherodactylus* to determine shared invasion patterns. To do this, we synthesized occurrence data to reconstruct the geographic and temporal invasion history and potential establishments of each of the *Eleutherodactylus* alien species. Using data from the native and likely established regions for each species, we assess whether realized climate niches are conserved between regions. Lastly, we evaluate the projected spread of these species under low and high greenhouse gas emission scenarios by using the native and established occurrence points to construct ecological niche models of future climatic conditions. This work provides insights into the abilities of species in the genus *Eleutherodactylus* to colonize regions with climatic conditions different from their native range, and projects their future spread for improved prevention and management of future introductions of these invaders.

## Methods

### Occurrence Data and Invasion History

Occurrence data for all five *Eleutherodactylus* species were downloaded on June 11th, 2025, from the Global Biodiversity Information Facility (GBIF.org 2025a-e). We ran an initial filtering step of all datapoints that contained geographic coordinates using the package *CoordinateCleaner* v 3.0.1 (Zizka et al. 2019) in R v 4.4.1 (R Core Team 2024), where we removed points with coordinates within the vicinity of country or province centroids, within the vicinity of known biodiversity institutions, as well as within the vicinity of known artificial hotspots from the Artificial Hotspot Occurrence Inventory (Park et al. 2023). Due to the small geographic regions these species have colonized within many countries (Lever 2003; Kraus 2009; Powell et al. 2011), we elected to not remove duplicate coordinates to ensure we did not miss biologically meaningful establishments within the same location. We also elected to include points without geo-referenced coordinates in the historic invasion reconstruction to avoid missing potential early introductions, but these points were later omitted in climatic niche comparisons. Furthermore, we removed 2 isolated records from the year 1860, including one from Peru for *E. antillensis* and one from Granada, Spain for *E. johnstonei*, that were well outside of the general invasive distribution for their respective species, and thus likely represented database entry errors. Using these filtered occurrence data, all countries for each species were classified into one of the following four categories: native, introduced, likely established, and artificial. Our criteria to assign records to non-native categories were based on the estimated generation time of all five *Eleutherodactylus* species in a previous study (3.00-3.46 years; Mancini et al. 2025). Populations were considered likely established when records were continuous in a country for at least three years, with no more than 3 years between observations, which suggests there has likely been more than one generation at that location. We considered a record as artificial if a specimen was recorded in an indoor environment or intercepted from a shipment by searching for the terms “greenhouse” and “shipment” in the location metadata. We combined records for many small, close-knit archipelago islands within the same country where migration between islands is likely high. For *E. johnstonei, E. martinicensis,* and *E. antillensis*, their native geographic origin has historically been ambiguous. We considered Montserrat, Antigua and Barbuda native for *E. johnstonei*, Martinique, Dominica, and Guadeloupe native for *E. martinicensis*, and Puerto Rico and the British Virgin Islands native for *E. antillensis* (see Supplementary Material 1 for a more detailed account on native ranges for the three species). We generated plots of occurrence records and their associated status using *ggPlot2* (Wickham et al. 2016).

### Climatic Niche Comparisons

To compare the realized niche of climatic conditions between native and invasive established regions for our five focal species, we restricted GBIF records to those that are considered native and likely established based on our three-year generation criteria. Using the *dplyr* package (Wickham et al. 2023), we further filtered data to only include the years 1951-2021 to match the corresponding climate raster data and also filtered points with a coordinate uncertainty of 5,000 meters or less, ensuring high accuracy when matching locality with climate data. Historic monthly weather data (1951-2021), specifically maximum temperature, minimum temperature, and precipitation, was downloaded from the WorldClim 2 database at a 2.5-minute resolution (Fick and Hijmans 2017).

We used the R package *raster* v 3.6-32 (Hijmans et al. 2025) to extract monthly values of maximum temperature, minimum temperature, and precipitation of the corresponding year for each datapoint. We then used the biovars function in the *dismo* package v 1.3-16 (Hijmans et al. 2020) to calculate all 19 bioclimatic variables from the extracted monthly minimum temperature, maximum temperature, and precipitation values. We used these 19 bioclimate variables to compare niche overlap between native and invasive climatic conditions for all five species using the package *ecospat* v 4.1.3 (Broennimann et al. 2012; Broennimann et al. 2026). Specifically, we used *ecospat* to run a principal component analysis of all 19 climate variables and obtained gridded climate values for each species’ realized niche across the first two principal components. While realized niches were gridded based on our native and likely established occurrence records, the background environment was established by obtaining bioclimate values of up to 10,000 random pseudoabsences based on the concatenated 1970-2000 bioclimate raster that was masked to landmasses using the Natural Earth database (South et al. 2024; Massicotte and South 2025). Randomized samples were chosen based on the minimum and maximum latitude and longitude from each native and invasive range. Pseudoabsences with missing climatic values were omitted. In the case of the restricted non-native range of *E. antillensis*, we were unable to extrapolate any pseudoabsences and thus considered the occurrence points reflective of both the realized and fundamental niche. From the gridded realized and fundamental niche spaces, we calculated Schoener’s D (Schoener 1968) and Hellinger’s I (Warren et al. 2008) overlap metrics. We also performed niche equivalency and similarity tests (Warren et al. 2008; Broennimann et al. 2012) using 1,000 total replicates for each test. For each respective test, we followed the COUE guidelines of invasive species niche quantification (Guisan et al. 2014) and explicitly assessed hypotheses of niche stability, or the proportion of grid space shared by native and invasive niche, niche expansion, or the proportion of unique grid space in the invasive niche, and niche unfilling, or the proportion of unique grid space for the native niche.

### Future Climatic Range Projections

To process occurrence points for future range projections, native and likely established datapoints from 1951-2025 outside of landmasses were removed using a global land polygon generated by combining the 10-minute-resolution “land” and “minor islands” polygons of the Natural Earth database (South et al. 2024; Massicotte and South 2025). Based on an estimated dispersal rate of approximately 1 km per year for *E. coqui* (Woolbright 1985), occurrence points were buffered by 40 km and 80 km and clipped to the global land polygon. The occurrence points were used to generate dispersal-constrained masking polygons for processing environmental layers for the periods 2041–2060 and 2081–2100, respectively, representing the maximum potential dispersal distances since 2020. Rarefying the occurrence points of each species was conducted by 1-km resolution (i.e., leaving one point per 1 km), subsequently generating a Gaussian kernel density with 0.04167 degree (ca. 2.5 arc-minutes, consistent with the climate data below) to account for sampling bias in downstream ENMs, and also to match with the resolution of the climate data used (see below). These steps were performed using SDMtoolbox Pro v 2.6 (Brown et al. 2017) and ArcGIS Pro™ v 3.4.3, following the SDMtoolbox developers’ recommendations.

In addition to the present (1951-2021) environmental data above, we downloaded future (2041-2060, 2081-2100) data from WorldClim using 2.5-minute resolution (Fick and Hijmans 2017). We chose two climate change scenarios: (i) low greenhouse gas (GHG) emission path with a 1.3–2.4 °C increase of global temperature (SSP 126, hereafter low GHG), and (ii) high greenhouse gas emission path with a 3.3–5.7 °C increase of global temperature (SSP 585, hereafter high GHG) from five global climate models (GCM), “ACCESS-CM2”, “INM-CM5-0”, “IPSL-CM6A-LR”, “MPI-ESM1-2-HR”, “UKESM1-0-LL” ensuring an adequate representation of varying climate projections from different modeling institutions. The present minimum monthly temperature, maximum monthly temperature, and monthly precipitation layer data were clipped using a 100-km buffer polygon created for each species. As in the *Climatic Niche Comparisons*, we calculated each of the 19 bioclimatic variables from the minimum temperature, maximum temperature, and precipitation using the ‘biovar’ function within the clipped extent. The 19 bioclimatic variables were summarized into PC axes according to the loading scores calculated from a PCA of all known established and native occurrence points used in the *ecospat* analysis. We used the PC axes explaining 95 % of the variance of the future environmental data for each species. We conducted all of these steps using ArcGIS Pro™ and RStudio 2023.12.0+369 (RStudio Team 2020) attaching R 4.3.2 (R Core Team 2021), and used the R packages *raster* v 3.6-32 (Hijmans 2024) or *terra* v 1.8-70 (Hijmans *et al*. 2024) for raster data handling and *sf* v 1.0-21 (Pebesma 2018) for geospatial vector data handling, respectively.

For the ENM algorithm, we chose MaxEnt (Phillips et al. 2006) based on its superior performance compared to other popular algorithms (Valavi et al. 2022). MaxEnt parameters were tuned using the R package *ENMeval* v 2.0.5.2 (Muscarella et al. 2014) for each species following the sampling of 10,000 or 50,000 random background points (Barbet-Massin et al. 2012; Valavi et al. 2022), generated with the *spatSample* function from the R package *terra*. These background points were sampled without replacement in proportion to sampling density derived from a previously generated kernel density layer, and the background sample size (10,000 or 50,000 points) yielding higher model performance was retained for the final ENM of each species later. To examine and control the complexity of the model response curves, a comprehensive set of feature classes [‘linear’, ‘linear and quadratic’, ‘linear, quadratic, and hinge’, ‘Hinge’] and regularization multipliers [0.5, 1.0, 1.5, 2.0, 2.5, 3.0, 3.5, 4.0, 4.5, 5.0] were tested using the 5-fold partition for computational efficiency. The optimal combination of parameters was determined by the Continuous Boyce Index (CBI) and used in the subsequent ENMs. We ran the ENMs using the R package *biomod2* v 4.2-6-2 (Thuiller et al. 2024). For *E. antillensis* and *E. martinicensis*, all available raster cells were used as background points in one set (n=351 and n=557 for *E. antillensis* and *E. martinicensis*, respectively) due to their limited range. Cross-validation (CV) of models was conducted with the five sets of randomly drawn CV points, with 80% of the data for calibration. We employed CBI, ‘Relative Operating Characteristic (ROC)’ and ‘True Skill Statistic (TSS)’ for the model evaluation (Miller 2010). After fitting individual models, an ensemble model was generated by computing a weighted mean of predicted probabilities across all individual models with CBI values greater than 0.5, using CBI as the weighting metric. The ensemble model was then projected to present and future conditions under low and high GHGs scenarios using each of the five GCMs. Finally, the models based on each GCM were combined into a single GCM ensemble model with equal weights.

Model validation statistics and ensemble variable importance values are included in Tables S1-S2.

## Results

### Historic Range Expansion

Our synthesis of occurrence records suggests that, despite the invasive success of each species, most introductions show no evidence of successful establishments (Figs. 1-3; Fig. S1). The earliest evidence for introductions in our dataset was from *E. johnstonei*, which was introduced into the United Kingdom and Grenada and *E. martinicensis* in Saint Kitts and Nevis Islands. Additionally, *E. planirostris* had multiple introductions into the United States, *E. johnstonei* on Saint Kitts and Nevis Islands, and *E. martinicensis* Trinidad and Tobago, and Saint Lucia, in the mid to late 1800s, that likely did not lead to establishments. Many introductions also occurred in the early 1900s, which also show no evidence of establishment. Those introductions that did not result in established populations include *E. johnstonei* in Barbados (with a brief likely establishment in the 1920s), Jamaica, Guyana, Saint Kitts, and Panama (Fig. 3), *E. planirostris* in Jamaica and the Dominican Republic (Fig. 2), *E. antillensis* on St. Croix, and *E. martinicensis* in Puerto Rico, Bermuda, St. Lucia, Antigua, Barbados, Jamaica, Grenada and Saint Vincent (Fig. S1).

**Fig. 1:**
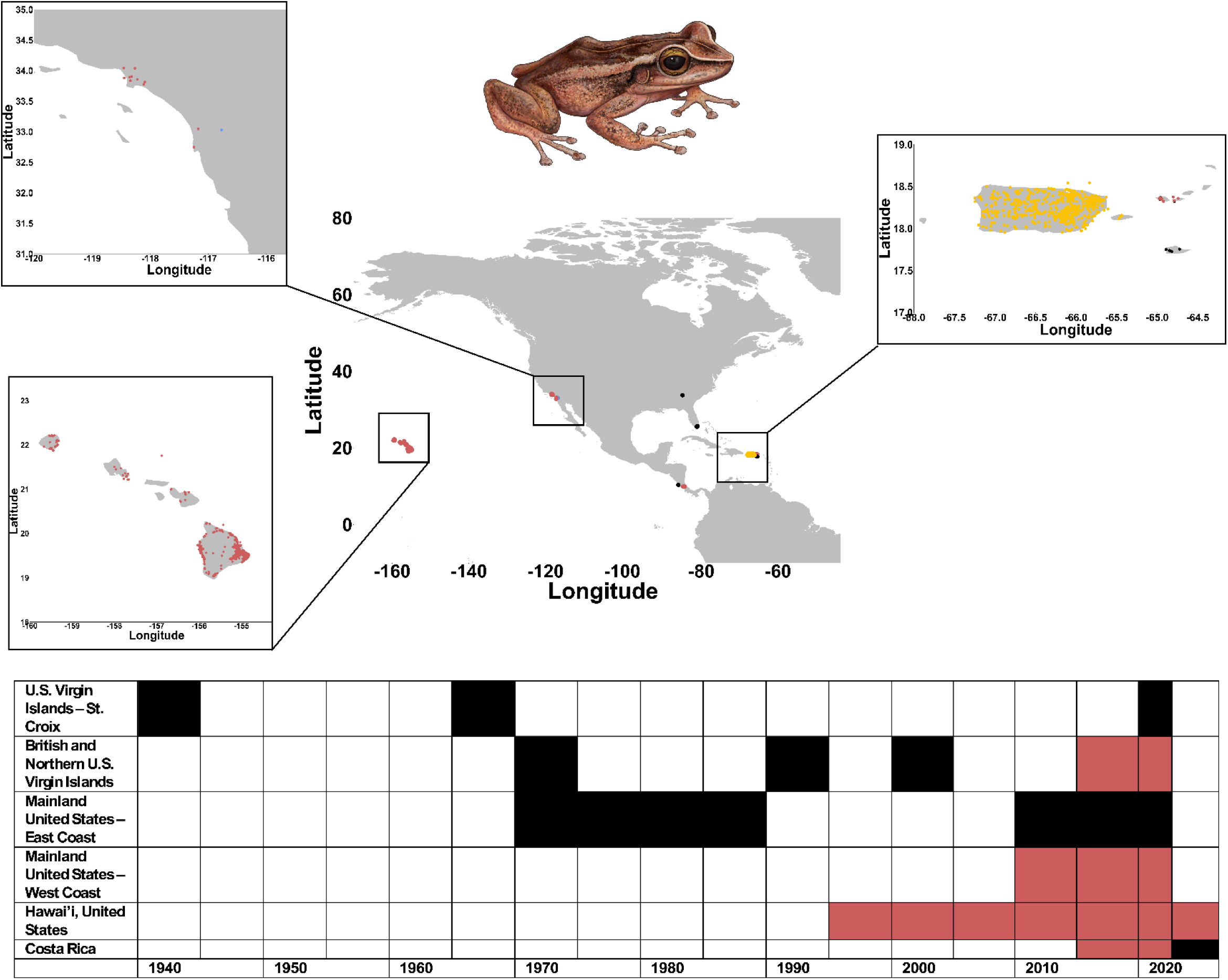
Location of native (yellow), likely established (red), and uncertain establishment (black) datapoints with known coordinates (top), and a timeline of historic introductions and establishments (bottom), for *Eleutherodactylus coqui*. Frog illustration by G. Sincich-Sosa.

**Fig. 2:**
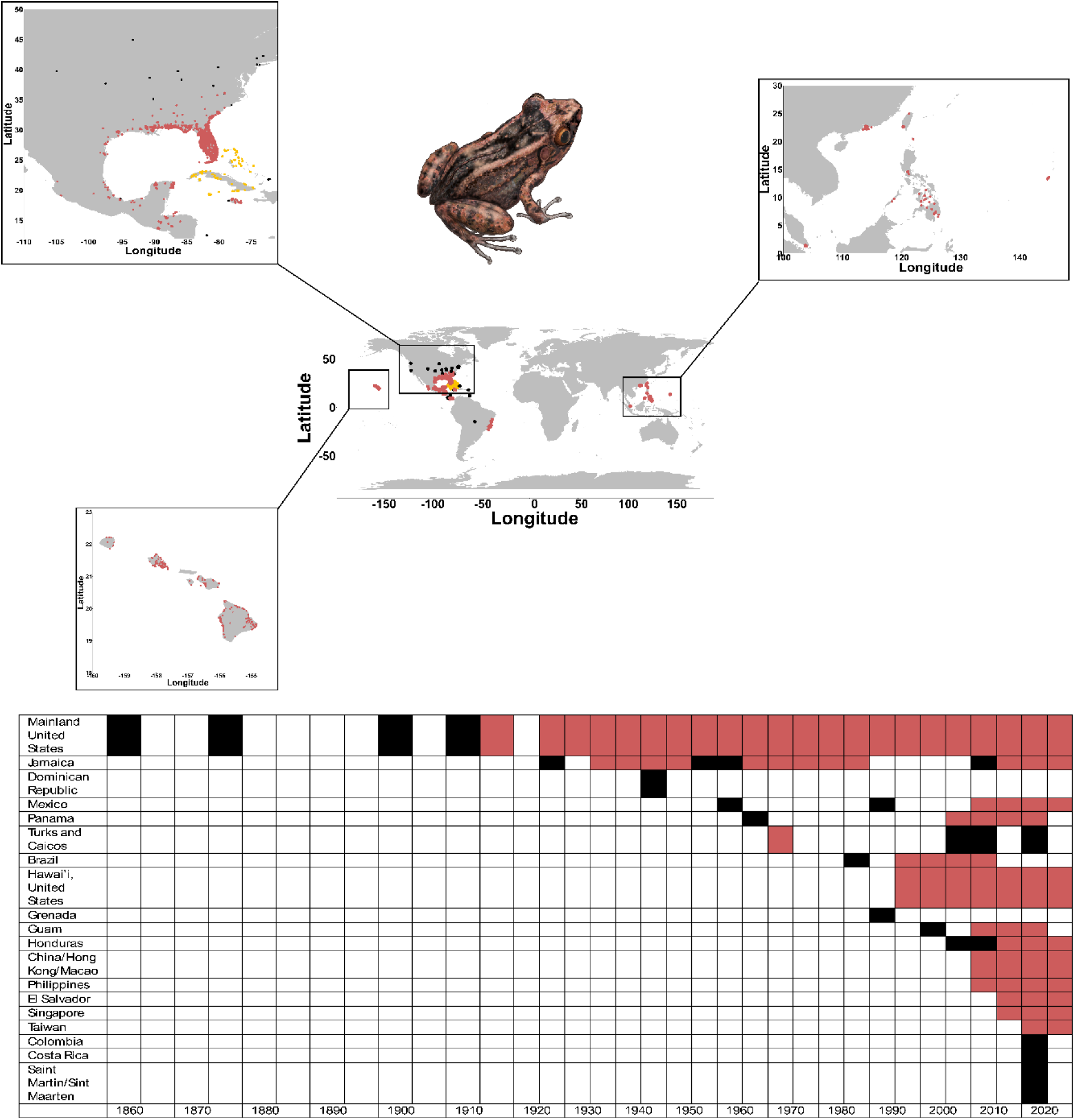
Map of the locations of native (yellow), likely established (red), and uncertain establishment (black) datapoints with known coordinates, and a timeline of historic introductions and establishments (bottom), for *Eleutherodactylus planirostris*. Frog illustration by G. Sincich-Sosa.

**Fig. 3:**
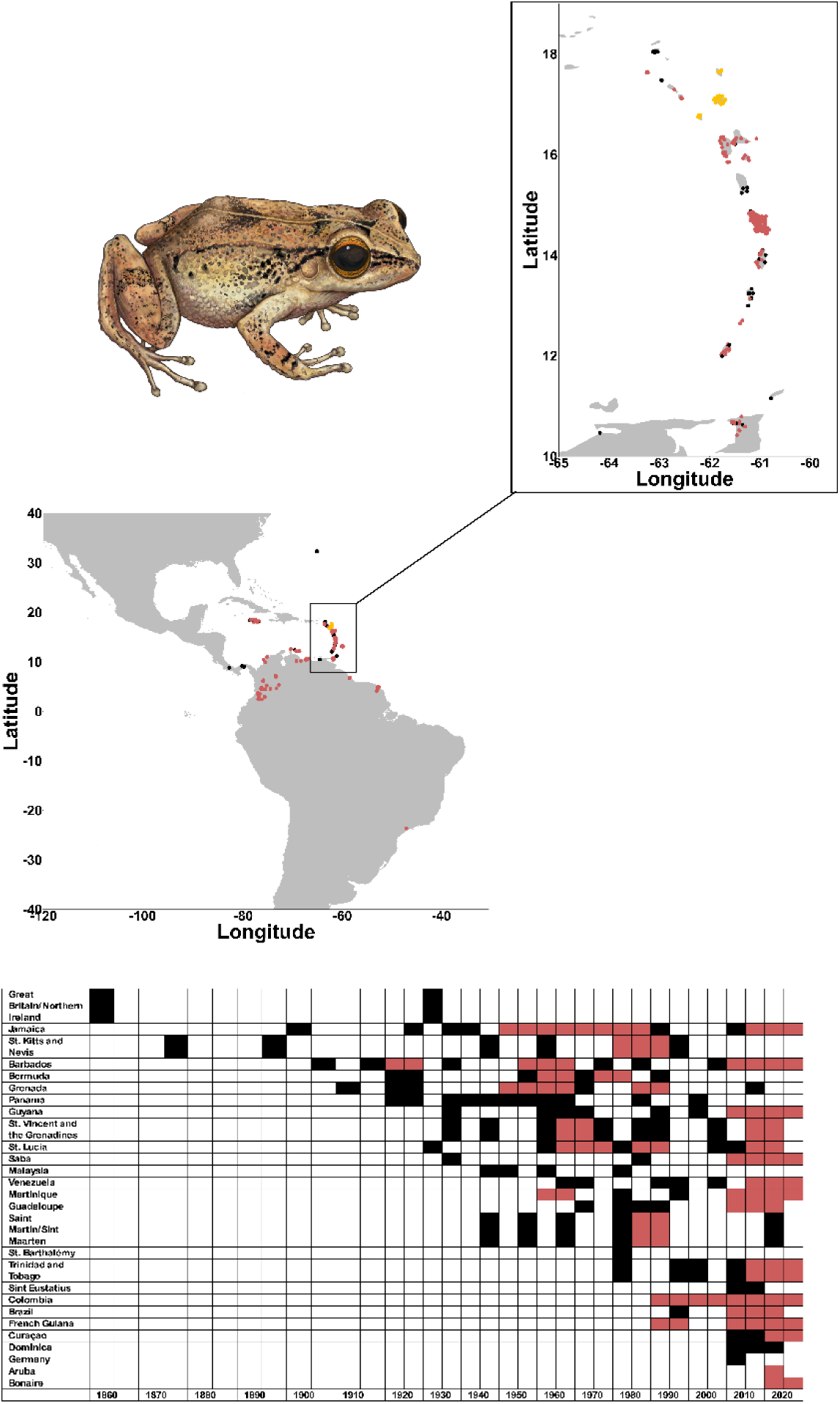
Map of the locations of native (yellow), likely established (red), and uncertain establishment (black) datapoints with known coordinates, and a timeline of historic introductions and establishments (bottom), for *Eleutherodactylus johnstonei*. Frog illustration by G. Sincich-Sosa.

The first likely successful establishment of any of these five species occurred when *E. planirostris* was introduced in the USA in the mid to late 1910s, where a population became established in Florida and has persisted to this day (Fig. 2). This species has also likely established in many countries and regions, including Jamaica, China, Mexico, Guam, Honduras, El Salvador, Brazil, and the Philippines, making it the most widespread invasive *Eleutherodactylus*. The first likely establishment of *E. johnstonei* occurred in Barbados in the 1920s, followed by likely establishments in Grenada and Jamaica in the early 1950s, but there is no evidence of long-term persistence of these populations (Fig. 3). Despite an early lack of persistent establishment, *E. johnstonei* is currently likely established in 14 countries, making it the most successful *Eleutheroactylus* in terms of the number of countries invaded. Establishment success for *E. coqui* occurred much more recently than the prior two species, with the earliest likely establishment occurring in Hawai’i in 1997. This species has also most recently become established in the British Virgin Islands and California (Fig. 1), but it remains to be seen whether these likely establishments persist. In contrast, *E. antillensis* and *E. martinicensis* have much reduced geographic reach and have become established in a more limited number of countries than the prior three *Eleutherodactylus*. While *E. antillensis* became established only on the U.S. Virgin Island of St. Croix, *E. martinicensis* was only established briefly in Jamaica and Grenada with no evidence of long-term persistence (Fig. S1).

### Climatic Niche Comparisons

All of our focal species showed evidence of different niche profiles between native and invasive ranges (Fig. 4; Table 1). *Eleutherodactylus martinicensis* showed the highest degree of overlap with a Schoener’s D and Hellinger’s I value of 0.34 and 0.47, respectively (Table 1), indicating a moderate degree of niche overlap. All other species had a low degree of niche overlap. Except for *E. antillensis*, all focal species showed evidence of a broader realized and background niche in their invasive range (Fig. 4). *Eleutherodactylus antillensis*, on the other hand, showed a broader native realized and fundamental niche. Despite these qualitative differences in niche breadth, the statistical evidence for niche expansion remains mixed. While niches between native and invasive ranges were similarly inequivalent for all taxa, only one species, *E. coqui*, showed evidence of significantly low niche similarity between the two ranges. Additional results highlight the complexity of niche comparisons as COUE tests for stability, expansion, and unfilling were all statistically significant for *E. planirostris*, *E. johnstonei* and *E. antillensis* (Table 1).

**Fig. 4:**
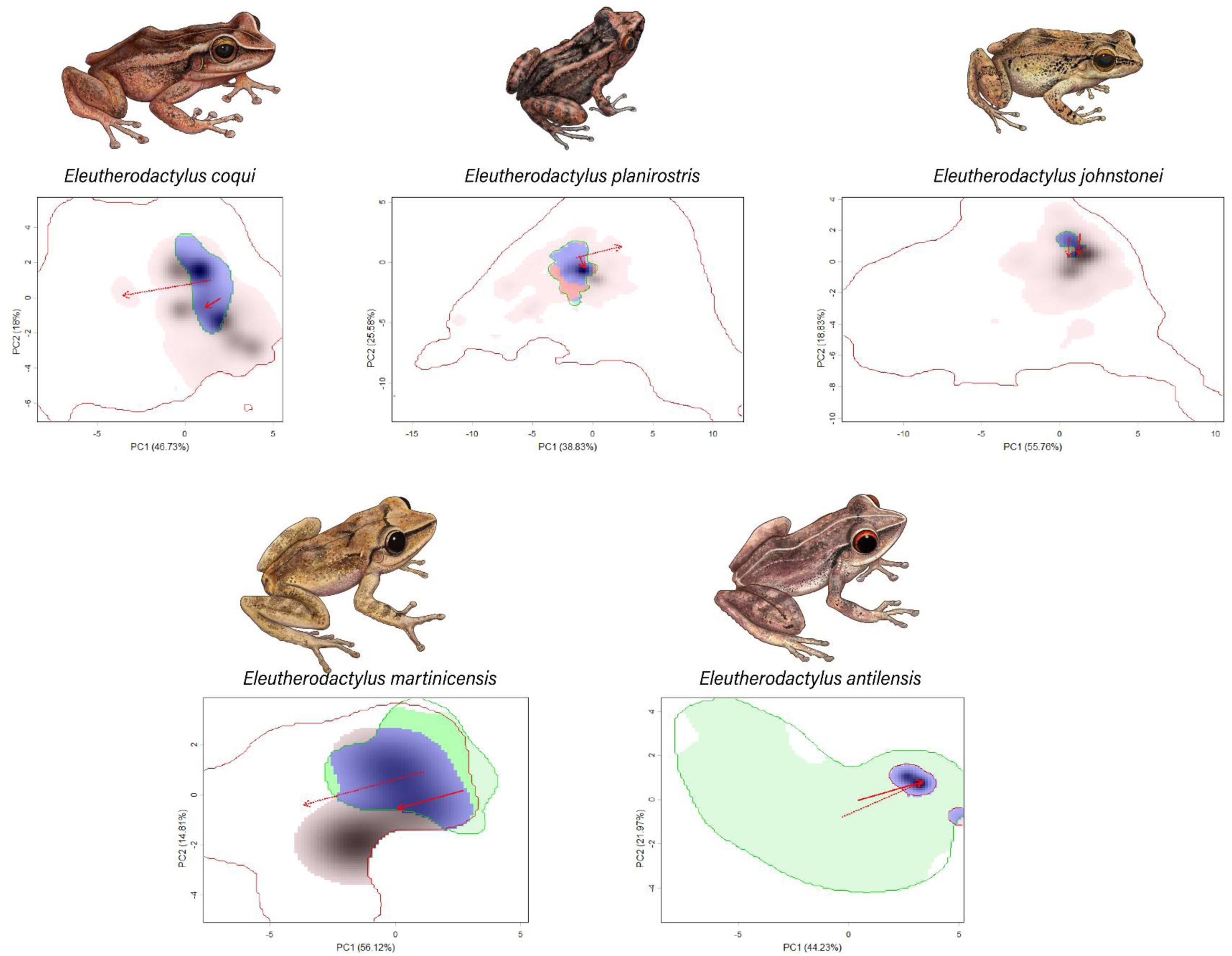
Ecospat analysis of climatic niches between native and invasive occurrence points for each invasive *Eleutherodactylus* species. Each figure showcases the native climatic niche (green), invasive climatic niche (red), their overlap (blue), and the direction of the niche shift (red arrow) for the first two principal component axes. Frog illustrations by G. Sincich-Sosa.

**Table 1:** Quantification of realized niche overlap between native and likely established invasive ranges for each *Eleutherodactylus* species. Statistics include Schoener’s D, Hellinger’s I, as well as statistical tests of Niche Equivalency and Niche Similarity, as well as similarity tests for Niche Stability, Expansion and Unfilling based on the COEUS framework.

|  | SCHOENER'S | HELLINGER'S | NICHE | NICHE | NICHE SIMILARITY |  |  |
| --- | --- | --- | --- | --- | --- | --- | --- |
|  | D | I | EQUIVALENCY | SIMILARITY | Stability | Expansion | Unfilling |
| *ELEUTHERODACTYLUS*<br>*COQUI* | 0.13 | 0.31 | 0.001 | 0.001 | 0.059 | 0.050 | 0.052 |
| *ELEUTHERODACTYLUS*<br>*PLANIROSTRIS* | 0.096 | 0.25 | 0.001 | 0.96 | 0.037 | 0.041 | 0.026 |
| *ELEUTHERODACTYLUS*<br>*JOHNSTONEI* | 0.046 | 0.15 | 0.001 | 1.0 | 0.001 | 0.001 | 0.001 |
| *ELEUTHERODACTYLUS*<br>*ANTILLENSIS* | 0.076 | 0.25 | 0.001 | 0.99 | 0.015 | 0.017 | 0.01 |
| *ELEUTHERODACTYLUS*<br>*MARTINICENSIS* | 0.34 | 0.47 | 0.001 | 0.92 | 0.10 | 0.094 | 0.077 |

Despite these mixed results, some commonalities in climatic niches can be drawn for our focal *Eleutherodactylus*. Our PCA analysis detected 3 variables – bio1, or the mean annual temperature, bio9, or mean temperature of the driest quarter, and bio11, or mean temperature of the coldest quarter – that showed consistently moderate to high PC loading scores (≥0.37) across both PC axes, indicating that these variables explain a high amount of variance in our climate dataset (Table S3). We illustrate these specific variables in Fig. 5, which shows statistically significant differences (Wilcoxon signed rank test, p ≤ 0.05) between native and likely established regions for each species, except for *E. coqui* for the mean temperature of the driest quarter between the two regions. These variables highlight common patterns of establishment into colder invasive regions for *E. coqui*, *E. planirostris*, and *E. johnstonei*. While we observe the opposite pattern for *E. antillensis* and *E. martinicensis*, with significantly higher temperatures in their invasive range, the climate conditions have a much narrower spread than the other three species, likely reflective of their restricted invasive range. It thus remains to be seen whether these two species can successfully establish in colder regions than those present in their native range.

**Fig. 5:**
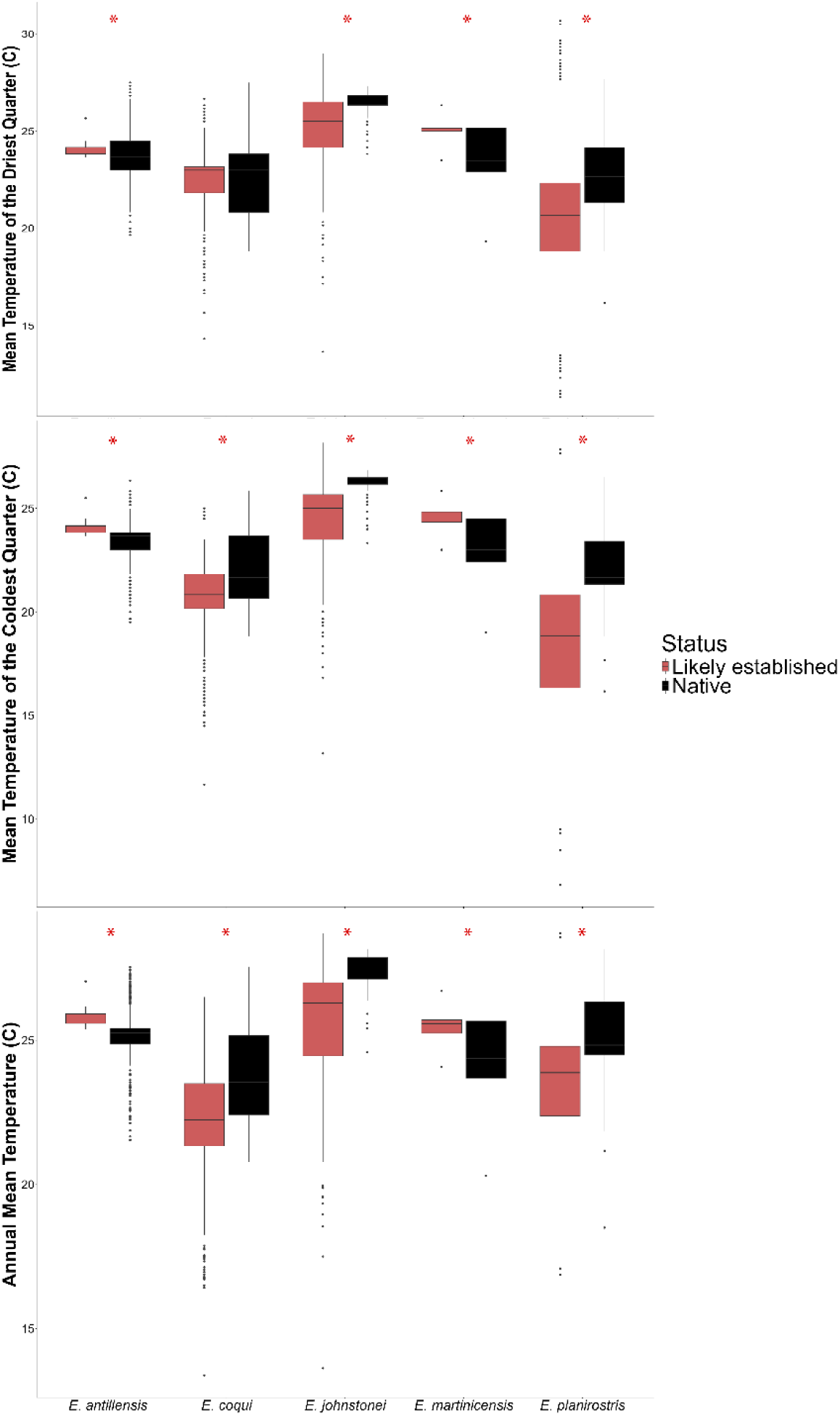
Comparisons of native and likely established niche space for all five invasive *Eleutherodactylus* species across the three variables with the highest principal component scores across all species. Red asterisk indicates a significant difference between native and likely established climate conditions based on Wilcoxon signed-rank test comparisons.

### Future Climate Projections

Maxent models of the current distribution of all five species indicate patchy suitable habitats largely centered around coastal regions (Fig. S2). Presently, *E. coqui* shows suitable habitat across the Big Island of Hawai’i, with higher suitability along the coast. Meanwhile, *E. planirostris* has the highest density of suitable habitat in the Southeastern US and also contains patchy but highly suitable habitat on many of the Hawai’ian Islands, as well as some patchy suitability in China, the Philippines, and the Yucatan Peninsula. In contrast to the large regions of high suitability for *E. coqui* and *E. planirostris*, *E. johnstonei* has a highly patchy distribution of suitable, smaller areas in South America, largely centered around cities, as well as the Lesser Antillean Islands. *Eleutherodactylus antillensis* and *E. martinicensis* have suitable habitats across the entirety of the islands where they have become established, including Jamaica (with a very low degree of suitability), Grenada and Martinique for *E. martinicensis* and Puerto Rico and the US Virgin Islands for *E. antillensis*. The northern portion of St. Lucia contains suitable habitats for *E. martinicensis*, a region for which there has not been evidence of establishment yet.

Under low GHG emissions, the suitable habitat of *E. coqui* on the Big Island of Hawai’i is largely stable, but under high GHG, portions of the interior of the island will increase in suitability, while the eastern coast has a marginal decrease in suitability (Fig. 6). In its native range, Puerto Rico, the suitable range is largely maintained under low GHG emissions, while the entirety of the island, especially the southern coast, is projected to be more unsuitable under high GHG emissions (Fig. S3). Similar to *E. coqui*, *E. planirostris* shows evidence of stable to very small increases in area in Hawai’i under low GHG emissions. However, under high GHG emissions, the suitability on the Big Island is expected to decrease, while largely remaining the same on the additional Hawai’ian islands (Fig. 7). Maintaining a patchy distribution of suitable habitats, *E. planirostris*’s habitat availability is expected to expand in Southeast Asia under both high and low GHG emissions, although most of the countries in this region have a low degree of suitability (Supplementary Figure 4). Increases in suitable habitat are also projected under both GHG scenarios in the United States and Caribbean Islands, with a contiguous pattern of high suitable habitat in the southeastern United States (Fig. 8). The suitable habitat projected for *E. johnstonei* is expected to marginally increase over its current range under both low and high GHG emissions, especially across the northern Andes and towards the Atlantic coast in Venezuela, but its established distribution in South America is likely to remain patchy (Fig. 9).

**Fig. 6:**
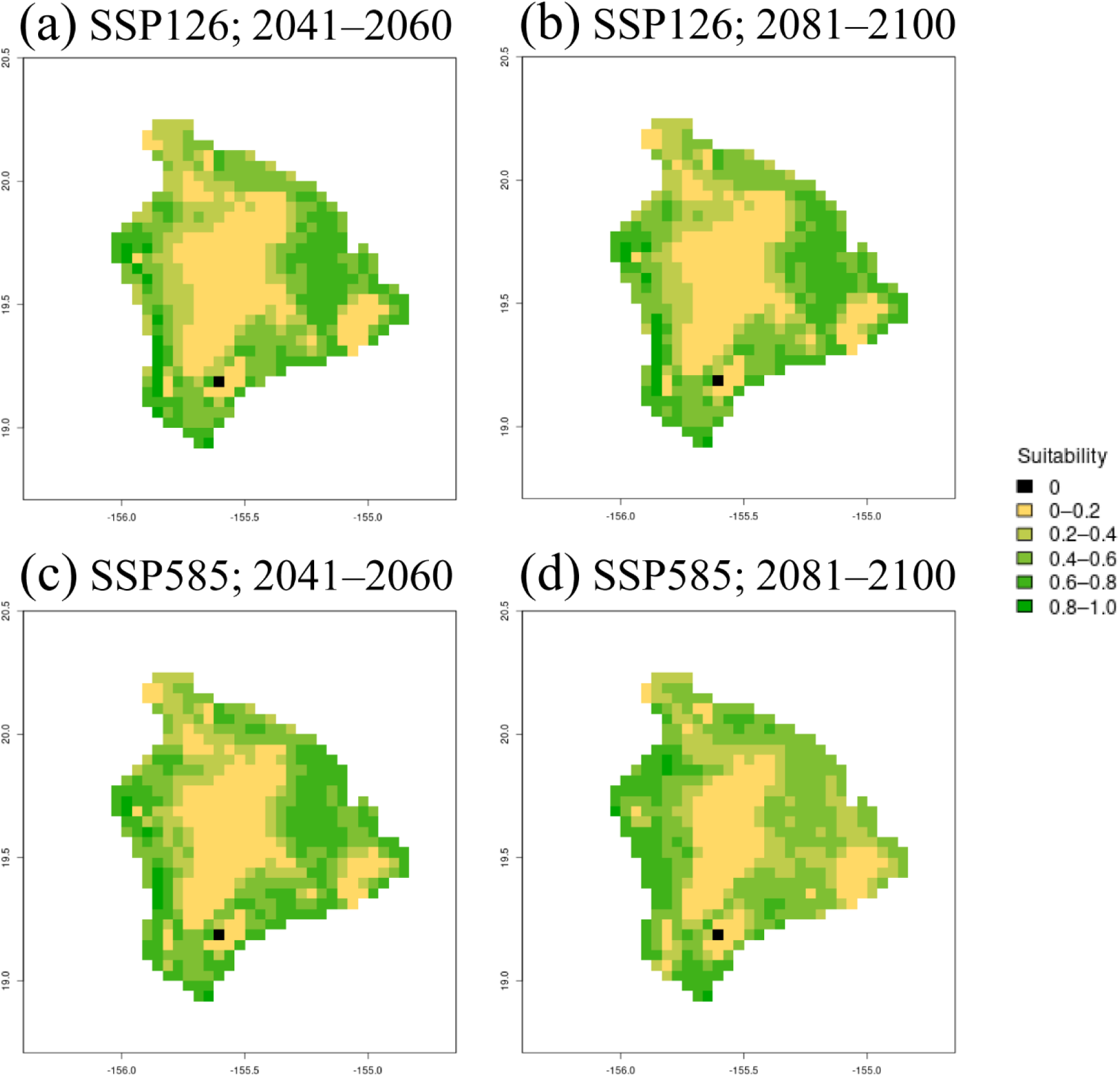
Projected invasive range of *Eleutherodactylus coqui* in the Big Island of Hawai’i under low (a, b) and high (c, d) GHG emission scenarios from 2041-2060 (a, c) and 2081-2100 (b, d). Black areas represent completely unreachable or unsuitable background land, while the other colors represent a scale from less (yellow) to more suitable (green) regions, respectively.

**Fig. 7:**
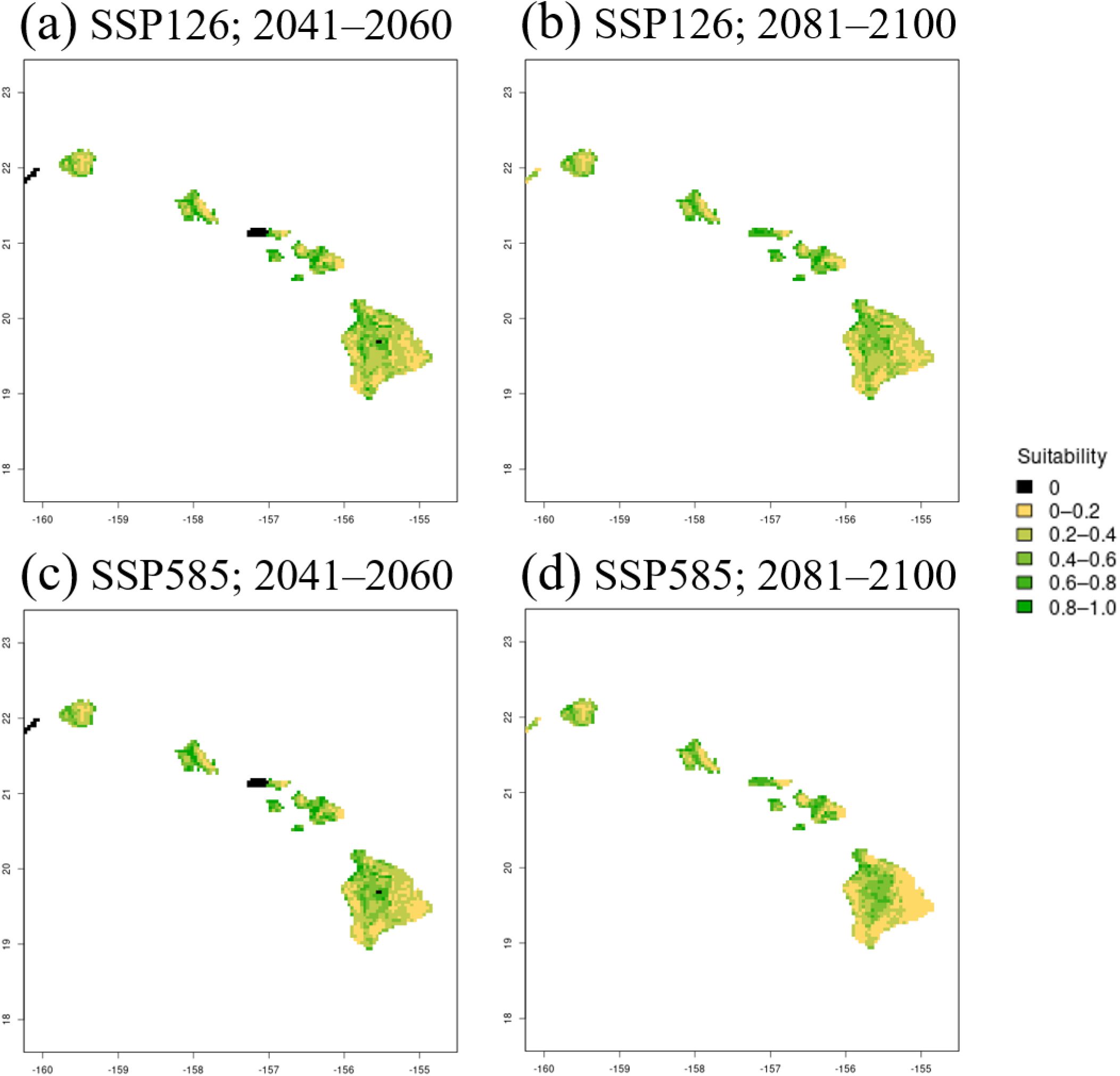
Projected invasive range of *Eleutherodactylus planirostris* in Hawai’i under low (a, b) and high (c, d) GHG emission scenarios from 2041-2060 (a, c) and 2081-2100 (b, d). The black landmass presents background land that is completely unreachable or unsuitable, while the scale of yellow and green landmass represents a scale of less and more suitable regions, respectively.

**Fig. 8:**
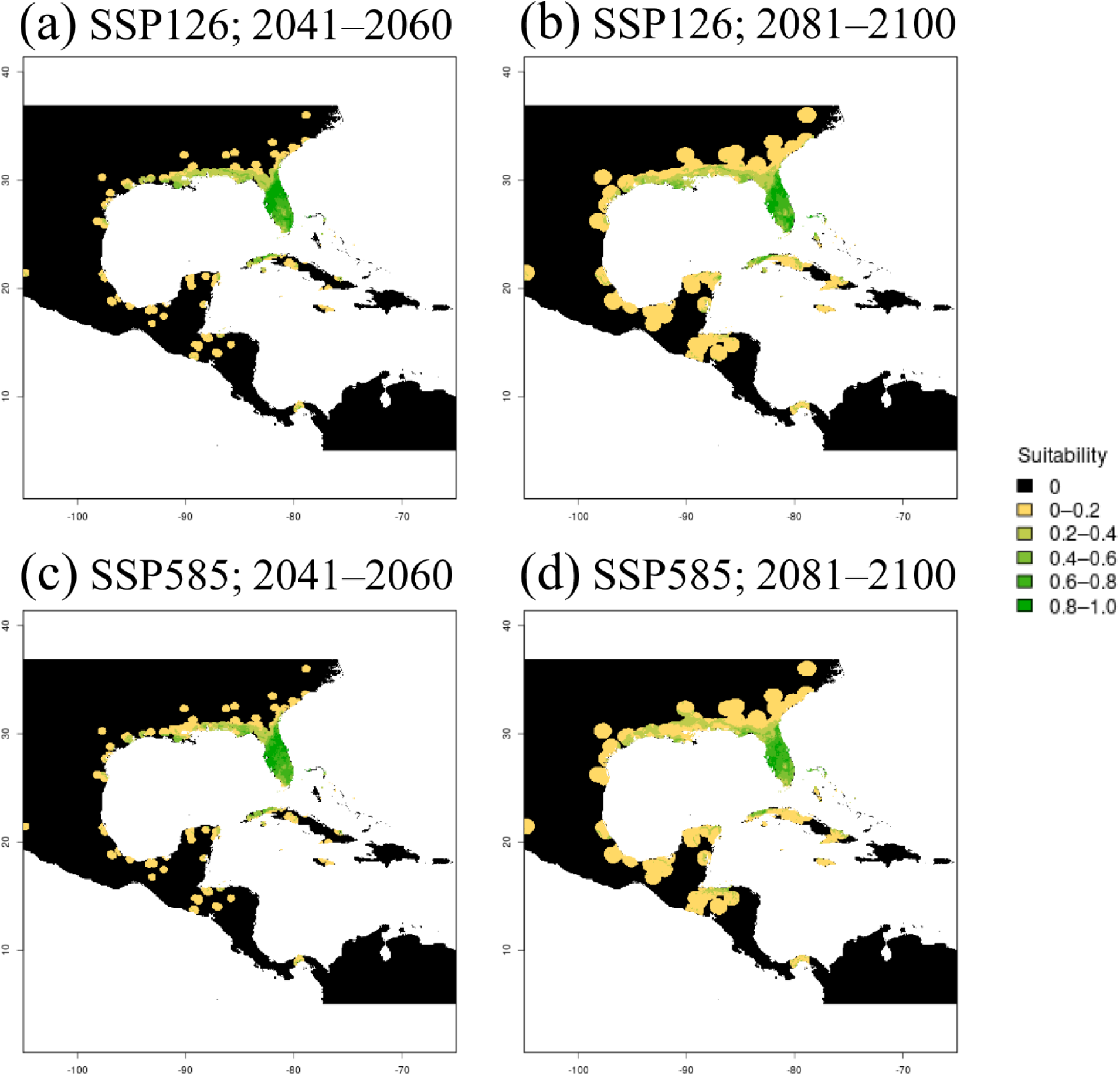
Projected invasive range of *Eleutherodactylus planirostris* in the North America and their native range of Cuba, Bahamas, and Virgin Islands under low (a, b) and high (c, d) GHG emission scenarios from 2041-2060 (a, c) and 2081-2100 (b, d). The black landmass presents background land that is completely unreachable or unsuitable, while the scale of yellow and green landmass represents a scale of less and more suitable regions, respectively.

**Fig. 9:**
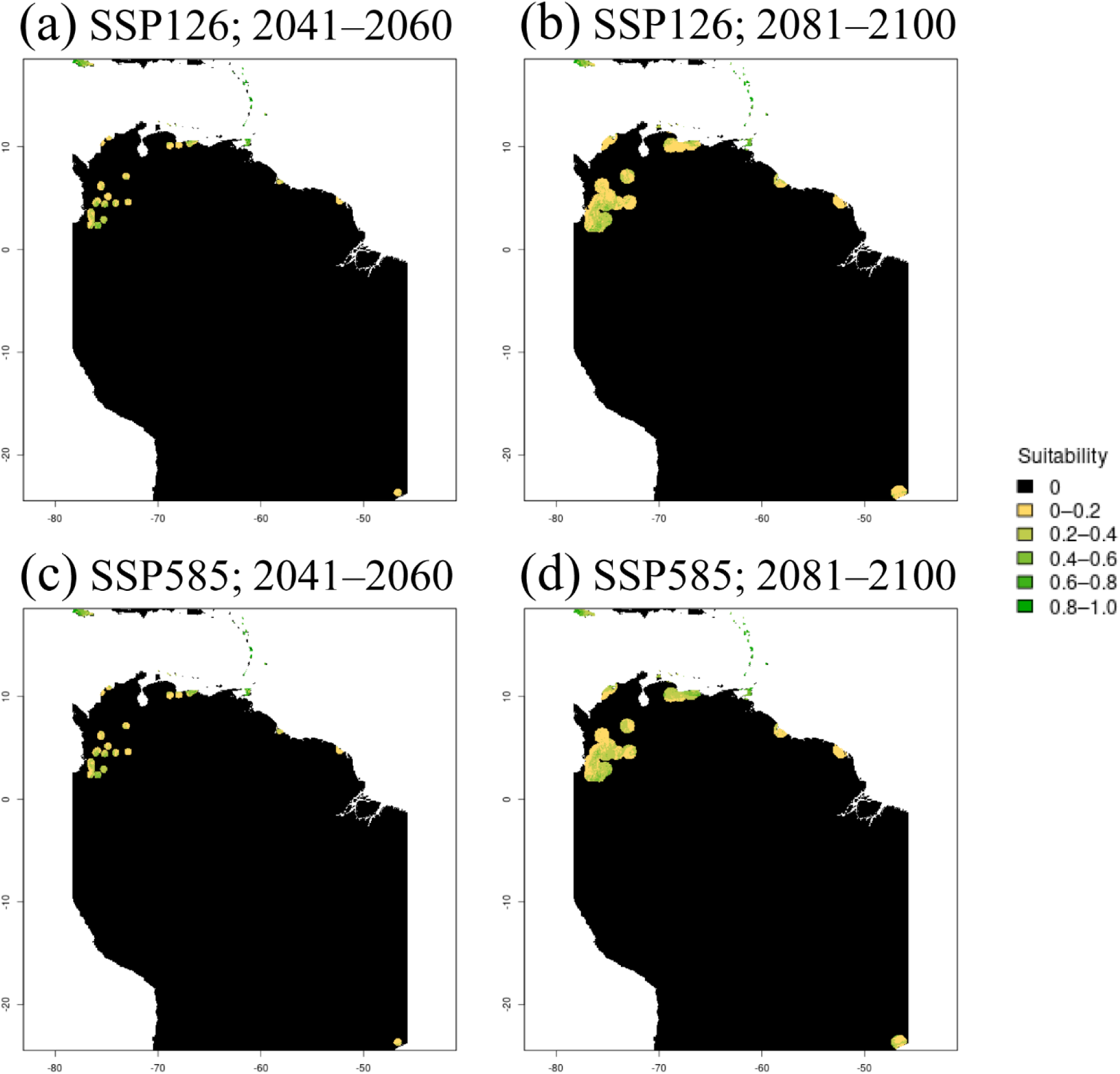
Projected invasive range of *Eleutherodactylus johnstonei* in South America and the Lesser Antilles under low (a, b) and high (c, d) GHG emission scenarios from 2041-2060 (a, c) and 2081-2100 (b, d). The black landmass presents background land that is completely unreachable or unsuitable, while scale of yellow and green landmass represents a scale of less and more suitable regions, respectively.

In comparison to the three most widespread *Eleutherodactylus* species, both *E. antillensis* and *E. martinicensis* have more restricted established distributions throughout their range and suitable habitat across most of the islands they inhabit. For *E. antillensis*, habitat suitability within their native Puerto Rico and invasive Virgin Islands is expected to remain stable under low GHG emissions and slightly decrease under high GHG emissions (Fig. S5), as was observed for *E. coqui* in Puerto Rico. The established populations of *E. martinicensis* within the Lesser Antillean Islands are expected to remain stable under low GHG emissions and potentially increase onto the island of St. Lucia where they have not yet become established (Fig. S6). Under high GHG emissions, certain regions in the Lesser Antilles, such as St. Lucia, will increase, while other islands, such as Martinique, will have small reductions in suitability. In Jamaica, *E. martinicensis* is projected to have a westward expansion in suitable habitat between the future time points under both emission (Fig. S7).

## Discussion

Comparing congeneric invasive species can reveal common patterns leading to their successful establishment in new geographic regions. In our case, our focal congeneric anurans show high variability in introduction history, and despite mixed statistical results, the three longest established and most successful invasive *Eleutherodactylus* – *E. coqui*, *E. planirostris,* and *E. johnstonei* - showed evidence of a broader invasive than native niche, with the ability to establish in colder regions likely driving these climatic niche shifts. In the future, suitable habitat for all five of our focal species is expected to remain stable, and even slightly increase, across regions under GHG emission scenarios, yet several projections showed slight regional-specific decreases in suitable habitat in later years (2081-2100) under high GHG scenarios.

For invasive alien species, it is frequently documented that they conserve their realized niche when introduced into new regions (e.g. Broennimann et al. 2012; Liu et al. 2020). How widespread such niche conservatism is in invasive alien species, however, is subject to debate (Bates and Bertelsmeier 2021) as they can also shift their realized climatic niches during biological invasions (Atwater et al. 2018). Such niche shifts have been documented in other anurans (e.g. Tingley et al. 2014; including *Eleutherodactylus* with mixed results – Li et al. 2014), likely due to their broad tolerance across an array of environmental conditions, a feature shared among many biological invaders (Sexton et al. 2002; Zerebecki and Sorte 2011). For our study, four of our five species showed a broader niche in their invasive range than their native range, with the three most widespread and earliest invasive species showing the largest differences between the native and invasive niche. For these analyses, we highlight that temperature, specifically the mean annual temperature and the temperature of the coldest and driest quarter, explains the largest degree of variance across all species. In addition, the three most widespread and oldest invasive species have become established in regions colder than their native range, suggesting a possible conserved strategy among these congeneric species to establish populations in these environmental conditions. This pattern is consistent with the idea that having a geographically restricted native range, with sufficient time to recover genetic diversity and acclimate or adapt to new environmental conditions, is likely to lead to climatic niche shifts (Alexander and Edwards 2010; Li et al. 2014). An exception to these climatic niche patterns is *E. antillensis* with broader environmental conditions in its native than in its established invasive range. However, this species is the most geographically restricted invasive *Eleutherodactylus* and is one of the more recently established invasive species. Thus, its sparse and recent introduction history likely explains its different climatic patterns when compared to other, more widespread *Eleutherodatylus*. Despite clear qualitative signals of niche expansion, we detected mixed results when implementing equivalency and similarity tests. Indeed, *E. coqui* is the only species showing statistically significant niche divergence under both tests. Under the COUE framework, three of our focal species, *E. planirostris*, *E. johnstonei,* and *E. antillensis,* show simultaneous evidence of niche stability, expansion, and unfilling. For these species, there is a significant degree of similarity between the invasive and native niche, yet the species also establish in climatic conditions differing from their native range and occur in invasive climatic regions absent in their current native distribution. These findings may be explained by a limited sample size or observer bias, discussed in more detail below, which can interfere with examining the role of chance on observed patterns of niche differentiation. Nevertheless, our results reveal that the species examined here shared their ability to establish in regions that are climatically unique from their native range.

There are many evolutionary mechanisms, including both phenotypic plasticity and adaptive responses, that likely explain the ability of *Eleutherodactylus* species to establish in novel climatic regions. Phenotypic plasticity is the first line of response of a species confronting novel environmental conditions (Lande 2016; Wang and Althoff 2019), and having high plasticity, specifically in thermal tolerance, may predispose species to become invasive (Sexton et al. 2002; Zerebecki and Sorte 2011). Such plastic responses to temperature have occurred in other invasive anurans such as cane toads in both Australia (McCann et al. 2018) and Florida (Mittan and Zamudio 2019), and painted reed frogs in South Africa (Davies et al. 2015). Our understanding of thermal plasticity in *Eleuthreodactylus* is still relatively sparse and has focused on a select few species, *E. coqui* and *E. antillensis*. These species have broader tolerance to extreme temperature conditions when compared to other, more distributionally restricted *Eleutherodactylus* (Beuchat et al. 1984; Delgado-Suazo and Burrowes 2022). However, other studies have shown that *E. coqui* was unable to acclimate to novel climatic conditions (Christian et al. 1988), suggesting that local adaptation, as opposed to plasticity, has played a stronger role in shaping the realized niche of this species. Plasticity in response to other abiotic factors may also promote the ability to become established in novel areas. For example, *E. johnstonei* can maintain steady oxygen consumption rates when humidity is reduced to 80% (Melo et al. 2026), which may support the species’ persistence during periods of low humidity when transported or establishing in a new region. However, overall, the role of plasticity remains understudied in the *Eleutherodactylus* invaders.

New environmental conditions can also impose strong selective pressures on invasive alien species, leading to rapid evolution. Evidence of this phenomenon, however, has historically been difficult to identify (Colautti and Lau 2015). Invasive alien species often undergo a population bottleneck, which reduces genetic diversity and adaptive potential (Nei et al. 1975). Nevertheless, evidence of rapid adaptation to new environmental conditions has been documented in introduced purple loosestrife (Colautti and Barrett 2013), pink salmon (Sparks et al. 2024), and cane toads (Tingley et al. 2012; Mittan and Zamudio 2019), to name a few examples. Consistent with founder events, non-native populations of *E. coqui* and *E. johnstonei* are depauperate of genetic variation (Peacock et al. 2009; Leonhardt et al. 2022). While such reduced genetic variation can limit adaptations to new conditions, it remains unclear whether these species have overcome the deleterious effects of low variation by deploying plastic or adaptive responses. Common garden experiments in *E. coqui* suggest plastic thermal responses are responsible for variance in body size, yet plasticity alone could not explain differences in body size variance between their native (Puerto Rico) and non-native (Hawai’i) distribution (O’Neill et al. 2018). Thus, more attention is needed to differentiate the influence of genetic and phenotypic variance in promoting successful establishment of *Eleutherodactylus* into new regions, and future studies should prioritize additional *Eleutherodactylus* species for genus-wide investigations of phenotypic plasticity and evidence of selection.

Despite differences in realized niche breadth between native and invasive ranges across the genus, habitat suitability is expected to remain stable or marginally increase for all five species under future climate change scenarios, but many habitats will likely be restricted and decrease in some regions under a high degree of GHG emissions. For *E. planirostris*, the suitable habitat is expected to increase under the low GHG emissions scenario along the coast of the Hawai’i, in the southeastern USA and Mexico, and the Philippines, but this species’ spread is more limited under high GHG emissions in many cases. Other studies projecting the climatic range of *E. planirostris* suggest a wider geographic spread (Mu et al. 2022), likely due to our conservative approach of including only ‘native’ and ‘likely established’ populations. By focusing only on those localities, our model reflects the finer-scale heterogeneity, and thus more precise estimates, of the different locations in which *E. planirostris* can potentially become established under future climate scenarios. *Eleutherodactylus coqui* and *E. johnstonei* also have the potential to expand their range into new regions, including the coast of the Big Island of Hawai’i for *E. coqui* and Colombia, Venezuela, Guyana, French Guiana, and Brazil for *E. johnstonei*. These range expansions largely corroborate previous studies (Rödder 2009a,b) yet are slightly more conservative in the estimated range area. Specifically, the predicted range expansion of *E. johnstonei* is restricted to major cities in northern South America where this species has not colonized non-urban areas (Gorzula 1989; Barrio-Amorós 2001; Kaiser and Henderson 1994; Leonhardt et al. 2019).

As our ecological niche modeling focuses on the regional climatic conditions associated with occurrence data, the effects of other relevant factors on the distribution and spread of *Eleutherodactylus* remain unassessed. Microhabitats, for instance, can allow species to become established in areas with apparent new abiotic conditions by avoiding exposure to potentially unfavorable environments (e.g. Kurtul et al. 2024). These climate refugia may explain how the *Eleutherodactylus* species investigated here occupy areas with general conditions different from those characteristics of their native range. Microhabitat use in *E. coqui* to reduce water loss during the dry seasons, for instance, is a common behavioral mechanism used to avoid deleterious environmental conditions (Beard et al. 2009). As predicted, the use of microhabitats with high relative humidity has been reported in the non-native ranges of some of these species. For example, *E. johnstonei* uses humid microhabitats consisting of leaf litter, grasses, rocks, and tree trunks in Bucaramanga, Colombia (Ortega et al. 2005a,b). The use of habitats containing a high density of captive-propagated plants, such as zoos, nurseries, or botanical gardens, also likely provides microhabitats expanding their range. *Eleutherodactylus* species may be present in artificial habitats, given that they are commonly introduced to greenhouses or nurseries. Some of the *E. coqui* records in California, for instance, were found in indoor greenhouses or were intercepted from plant shipments, and there are published records of *E. johnstonei* in the Czech Republic from a greenhouse in Prague (Moravec et al. 2020). While we filtered out the few occurrence points showing evidence of the individual being intercepted in an indoor greenhouse or in plant shipments, more complete documentation records of artificial settings are necessary to confidently assign the context in which all individuals were found.

Occurrence records across the native and invasive range of our focal species reveal discrepancies in niche expansion among *Eleutherodactylus* species. It is unclear, however, whether such differences reflect differences in detection rates and sampling bias among species. Differences in their call properties, for instance, have resulted in varying success in identifying and eradicating invasive species. In Hawai’i, *E. coqui* and *E. planirostris* are both present, yet the louder call of *E. coqui* leads to a higher detection probability and potentially masks the detection of the quieter *E. planirostris* (Olson et al. 2012). In Guam, *E. coqui* was quickly detected based on its loud call and eliminated soon after its introduction while *E. planirostris* successfully became established (Christy et al. 2007a). In addition to detection bias, sampling bias is well known to affect niche modelling (Barber et al. 2022) and can come from a variety of sources including bias for certain taxonomic groups (Reddy and Dávalos 2003) or toward geographic regions frequented by humans (Kadmon et al. 2004). While we have taken multiple steps to minimize sampling bias in our data with our filtering and processing steps, it is impossible to completely eliminate such bias, especially when considering early species occurrence records due to limitations in species detection and data sharing efforts. However, given that the spread of *Eleutherodactylus* is primarily through human trade (Christy et al. 2007a,b; Kraus 2009; Powell et al. 2011), concerns about detection are probably minor. Future work should prioritize systematic surveys of each species’ non-native range to identify breeding populations. Similarly, comprehensive population genomic analyses would be valuable to evaluate hidden pathways of introduction.

Overall, our study reveals unique, species-specific introduction histories and pathways with limited but general commonalities in establishment success among invasive alien *Eleutherodactylus*. Despite mixed statistical evidence for niche differentiation, most of our focal species show some degree of divergence between native and invasive niches. The majority of our species display the ability to establish in colder conditions, suggesting this pattern is a potentially conserved trait. Additionally, all five species are expected to continue their range expansion, to a limited degree, under a low level of GHG emissions but may experience some contractions under high GHG scenarios. This study suggests that abiotic conditions of the native range in *Eleutheroadactylus* are not necessarily indicative of the environment of the locations where they can become established, and regions with climates colder than in the Caribbean should not be assumed to be unsuitable for these invasive species. Thus, continued monitoring and the immediate capture of these invasive species, in addition to recommended nursery plant treatments to prevent their translocation to new areas (e.g. hot water, concentrated caffeine, or citric acid sprays, Global Invasive Species Database, 2025), should be implemented to prevent future establishments. Overall, our study suggests that tolerance to broad climatic conditions, whether through physiological plasticity, evolutionary adaptation, or microhabitat use, are likely factors making these *Eleutherodactylus* species highly successful invaders.

## Acknowledgements

We would like to thank members of the Bernal Lab for feedback on earlier versions of this manuscript. We are particularly grateful to Desi M. Joseph, who shared with us the pipeline for GIS processing of climatic data, and to Addy E. Messerly, who helped with data curation. We would also like to thank Gabriela Sinich-Sosa for creating the illustrations of three of the *Eleutherodactylus* species included in this study.

## Funding

AJM and XEB were supported by the National Science Foundation IOS-2054636.

## Competing Interests

The authors declare no competing interests.

## Author Contributions

AJM, JTK and XEB conceived the study. AJM and JTK ran initial preliminary analyses. AJM and JYJ developed the final analytical pipelines and analyzed the final data. All authors contributed writing, revising and approved the final version of the manuscript.

## Data Availability

Manually curated occurrence data are included in the Supplementary Material 2. Scripts used in data processing and analyses are archived in GitHub at https://github.com/andrewmularo/eleutherodactylus_spatialecology

## Notes

### Competing Interest Statement

The authors have declared no competing interest.

