## Supplementary Material 1 for "Disparate introduction histories but similar climatic distribution patterns among congeneric invasive anurans"

**Uncertain native origins of *Eleutherodactylus***

Many species within the genus *Eleutherodactylus* have historically been misidentified and records of endemism are poorly documented. For example, it was originally thought that *E. antillensis* was endemic to St. Croix (Lever 2003). However, later records indicate that St. Croix is part of the invasive range (Kraus 2009; Powell 2011), which has been supported by genetic studies (Barker et al. 2012; Barker and Rodriguez-Robles 2017). The IUCN Global Invasive Species Database (GISD) considers the US Virgin Islands as part of the invasive range of *E. antillensis* (Global Invasive Species Database 2025a). For this study, we consider the native range of *E. antillensis* to consist of Puerto Rico and the British Virgin Islands. *Eleutherodactylus martinicensis* has had similar problems with uncertain endemism, although there has been agreement that Martinique and Dominica are part of the native range (Schwartz 1961; Censky and Kaiser 1999; Kaiser and Hardy 1994; Lever 2003; Powell et al. 2011). Most of these early accounts consider Guadeloupe as part of their native range, yet recent records from the IUCN list Guadeloupe as part of the invasive range (IUCN SSC Amphibian Specialist Group 2021a; Global Invasive Species Database 2025b). Despite being listed, we find no compelling evidence with the ISSG or the IUCN red list that Guadeloupe represents an introduced population of *E. martinicensis*, suggesting that they were "probably introduced" (IUCN SSC Amphibian Specialist Group 2021a). Thus, we consider the native range of this species to consist of Martinique, Dominica and Guadeloupe.

Out of all our included species, however, none have had more disagreement in endemism than *Eleutherodactylus johnstonei*. Historically, the species has been confused with *E. martinicensis* (Kraus 2009) and occurs on virtually every island of the Lesser Antilles. The earliest accounts considered *E. johnstonei* to be endemic to the following islands: St. Vincent, St.

Lucia, Martinique, Antigua, Montserrat, Nevis, St. Eustatus, Saba and St. Martin (Schwartz 1961; Kaiser 1997). These accounts contained differing opinions on whether *E. johnstonei* was native to Barbados, Barbuda, Grenada, St. Christopher, and St. Kitts. Later accounts considered all of these islands, as well as St. Lucia, St. Vincent, Martinique, St. Eustatus and St. Martin as part of the invasive range (Kraus 2009; Powell 2011). Recent genetics studies have attempted to uncover the endemic region of *E. johnstonei*, with Yuan et al. (2022) suggesting Montserrat as the origin population, and Leonhardt et al. (2022) considering St. Lucia as the native origin. Currently, the IUCN considers Antigua and Barbuda as the likely native origin (IUCN SSC Amphibian Specialist Group 2021b), and the GISD considers St. Lucia as a part of its invasive range (Global Invasive Species Database 2025c).

For *E. johnstonei*, we consider Montserrat as a part of the native range based on genetic evidence from Yuan et al. (2022). Consistent with the IUCN, we also consider Antigua and Barbuda as a part of the native range, despite some arguments against this conclusion. Yuan et al. (2022) claim that Antigua is an introduced population based on low genetic diversity, yet this population falls under a completely different genetic clade than the Montserrat population. Thus, Antigua cannot be definitively excluded as part of the native range based solely on having low genetic diversity. Although Kaiser (1997) considered Barbuda inhospitable to anurans until colonial establishment, fossil records of anurans exist on the island (Lynch 1966), and Yuan et al. (2022), who did not sample in Barbuda, suggest the region as a possible secondary origin of *E. johnstonei*. Despite arguments to include St. Lucia as a native region, much of the recent consensus considers the population to be non-native (Powell 2011; Kraus 2009; Yuan et al. 2022; IUCN SSC Amphibian Specialist Group 2021b; Global Invasive Species Database 2025c), and some genetic studies have only assumed, but not definitively confirmed, that St. Lucia is a native region (Leonhardt et al. 2022). More comprehensive genomic sampling across the Lesser Antilles will be needed to confirm the true endemism of *Eleutherodactylus johnstonei*.

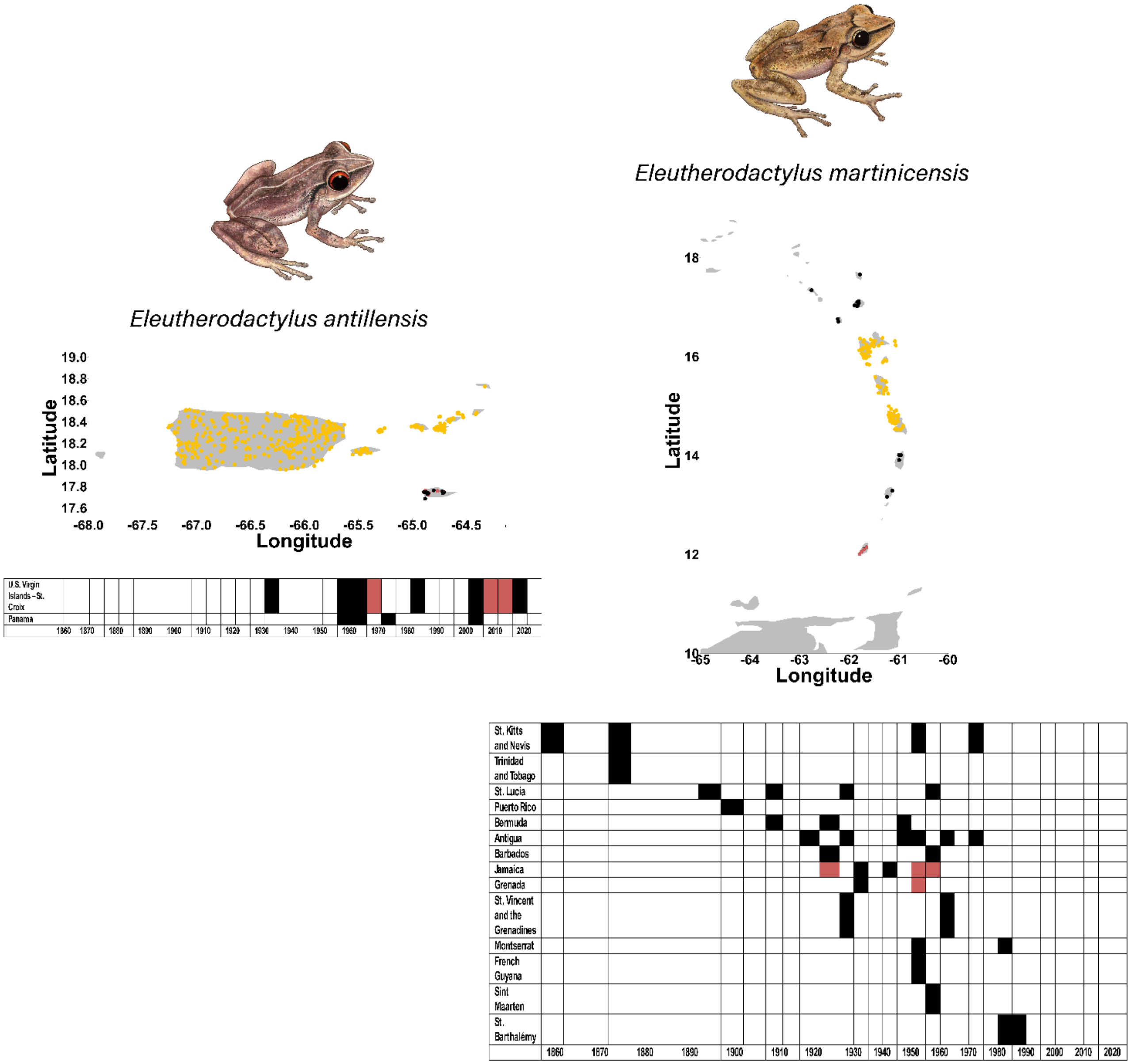
Eleutherodactylus antillensis
Latitude
Longitude
19.0
18.8
18.6
18.4
18.2
18.0
17.8
17.6
-68.0
-67.5
-67.0
-66.5
-66.0
-65.5
-65.0
-64.5
U.S. Virgin Islands – St. Croix
Panama
1860 1870 1880 1890 1900 1910 1920 1930 1940 1950 1960 1970 1980 1990 2000 2010 2020
Eleutherodactylus martinicensis
Latitude
Longitude
18
16
14
12
10
-65
-64
-63
-62
-61
-60
St. Kitts and Nevis
Trinidad and Tobago
St. Lucia
Puerto Rico
Bermuda
Antigua
Barbados
Jamaica
Grenada
St. Vincent and the Grenadines
Montserrat
French Guyana
Sint Maarten
St. Barthalémy
1860 1870 1880 1890 1900 1910 1920 1930 1940 1950 1960 1970 1980 1990 2000 2010 2020

**Fig. S1:** Map of the locations of native (yellow), likely established (red) and uncertain establishment (black) datapoints with known coordinates, and a timeline of historic introductions and establishments, for *Eleutherodactylus antillensis* and *E. martinicensis*.

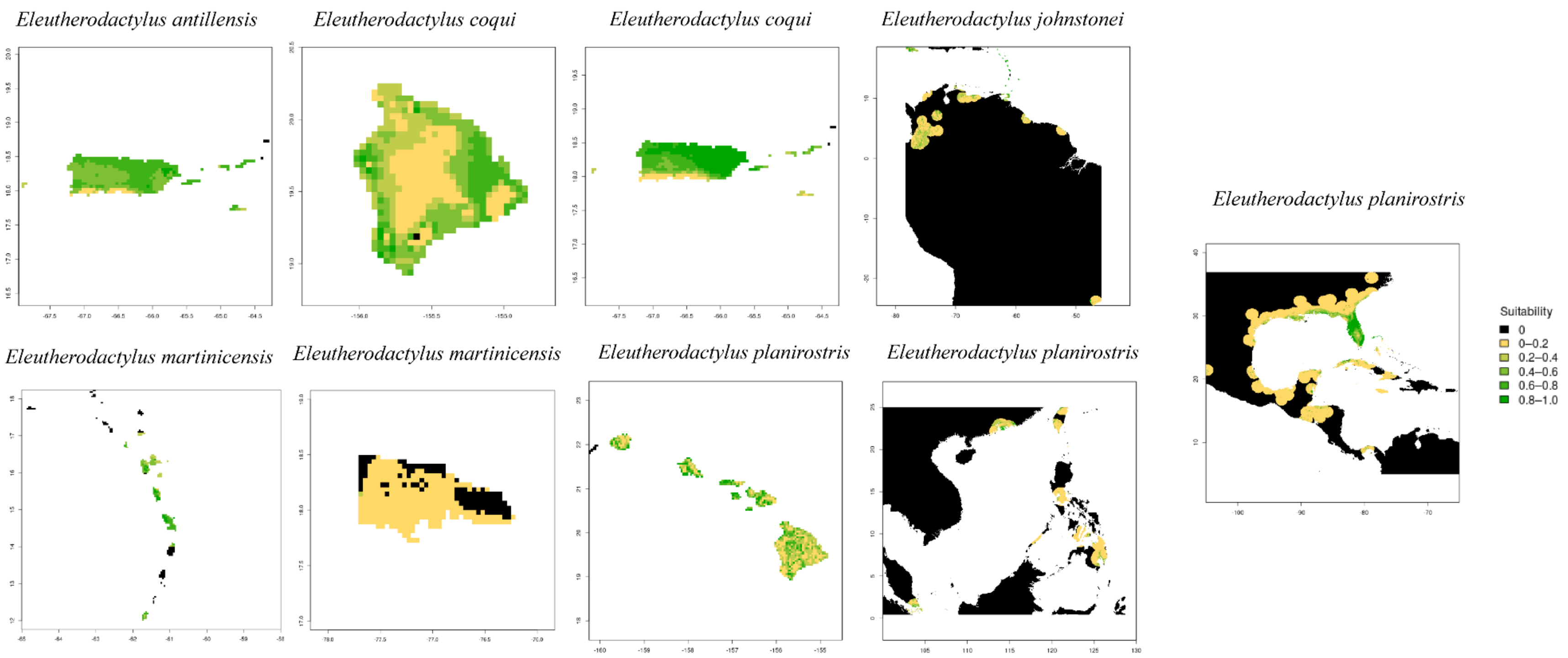


**Fig. S2**: Maxent models of the current distribution of each species. The black landmass presents background land that are completely unreachable or unsuitable, while scale of yellow and green landmass represents a scale of less and more suitable regions, respectively.

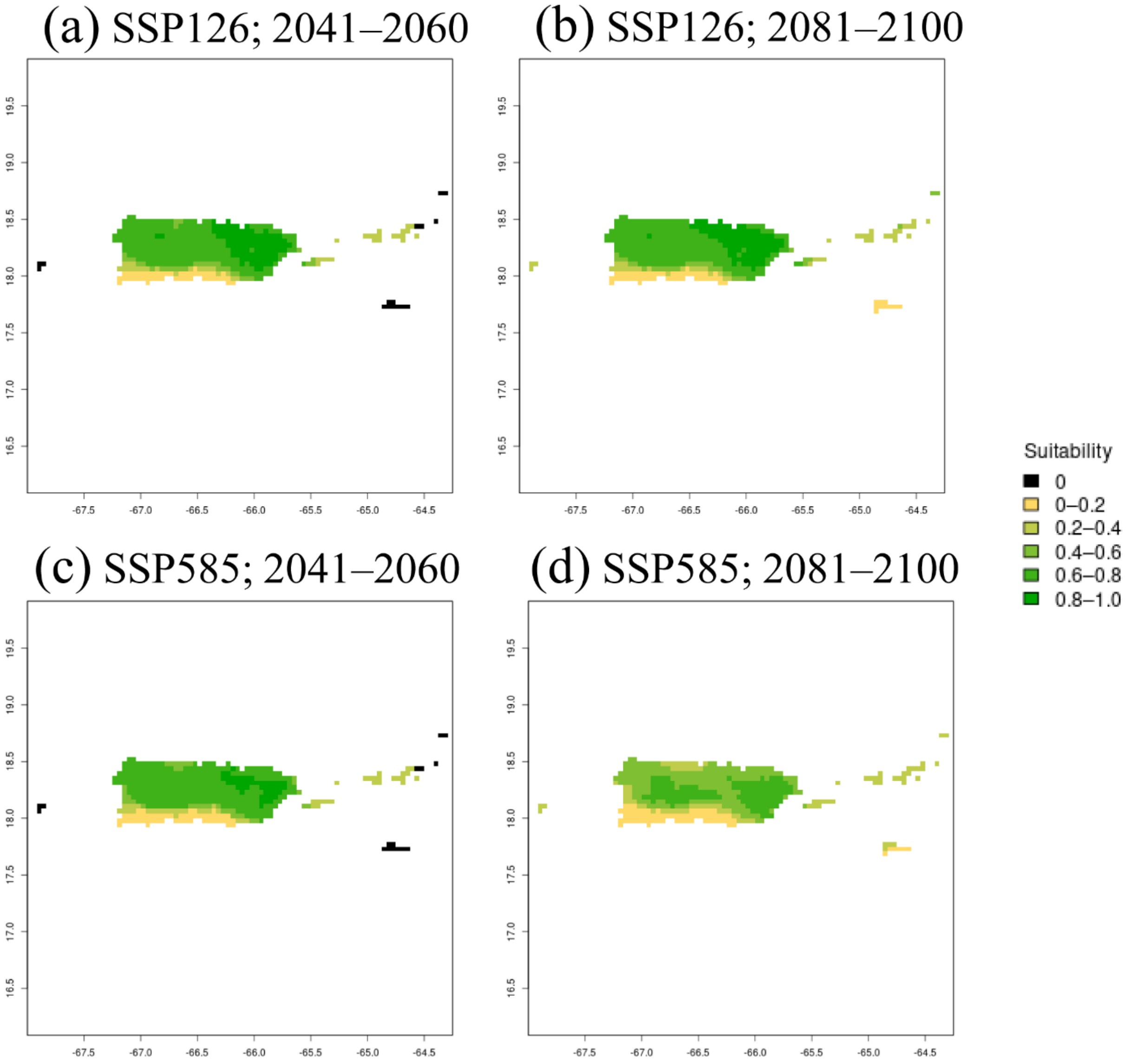


**Fig. S3**: Projected invasive range of *Eleutherodactylus coqui* in their native range of Puerto Rico under low (a, b) and high (c, d) GHG emission scenarios from 2041-2060 (a, c) and 2081-2100 (b, d). The black landmass presents background land that are completely unreachable or unsuitable, while scale of yellow and green landmass represents a scale of less and more suitable regions, respectively.

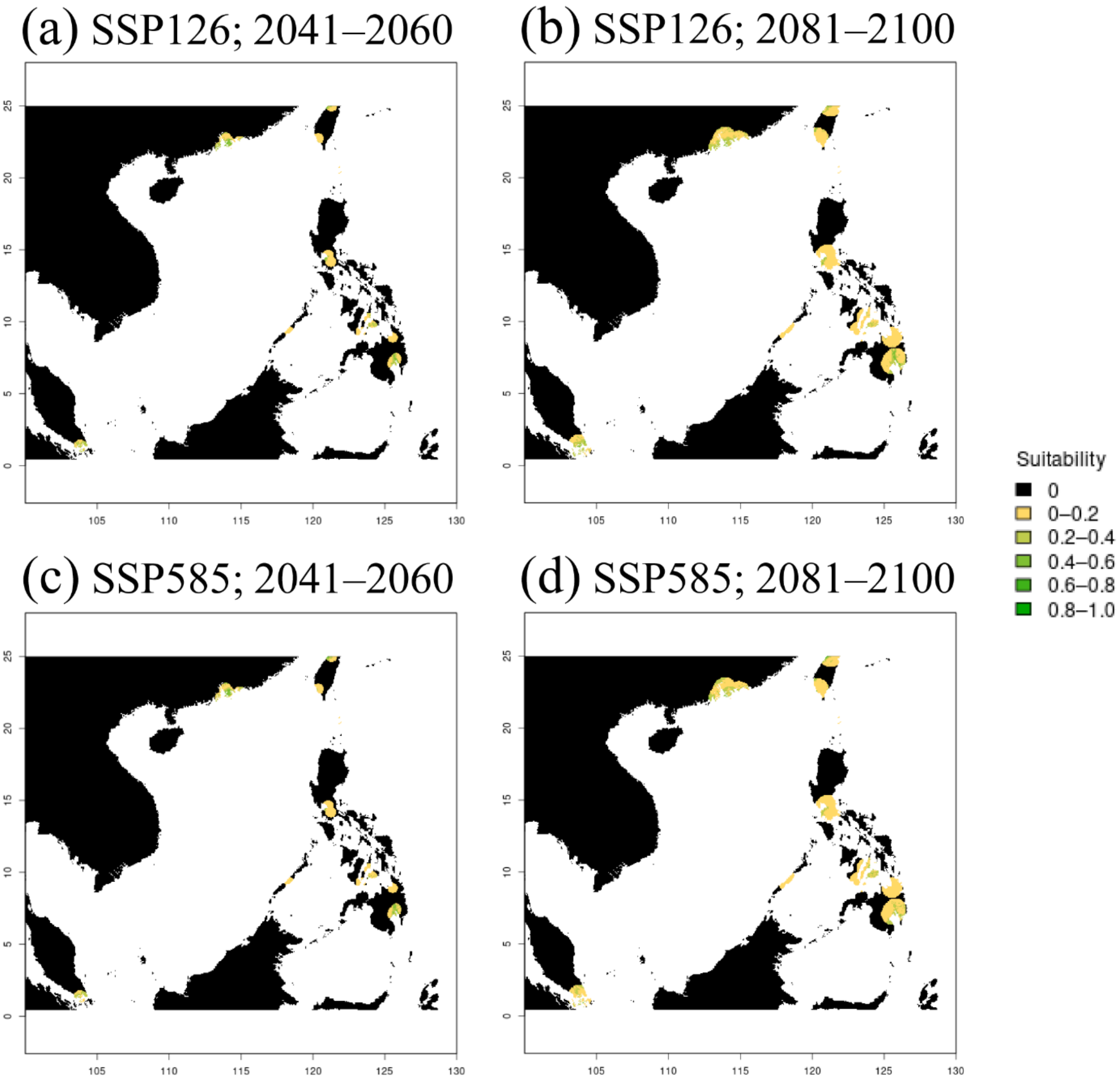


**Fig. S4**: Projected invasive range of *Eleutherodactylus planirostris* in Asia under low (a, b) and high (c, d) GHG emission scenarios from 2041-2060 (a, c) and 2081-2100 (b, d). The black landmass presents background land that is completely unreachable or unsuitable, while scale of yellow and green landmass represents a scale of less and more suitable regions, respectively.

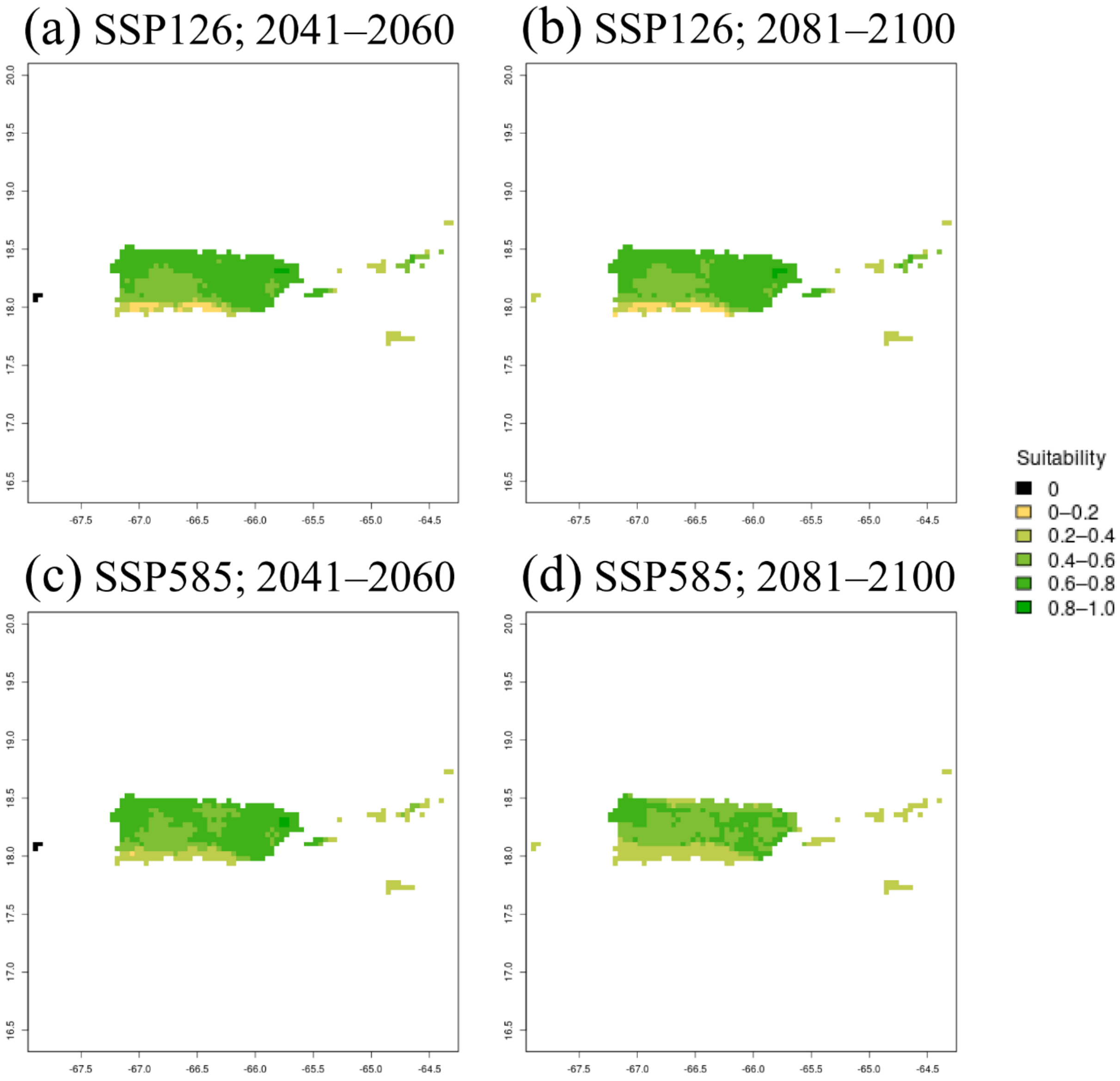


**Fig. S5**: Projected invasive range of *Eleutherodactylus antillensis* in Puerto Rico and the Virgin Islands under low (a, b) and high (c, d) GHG emission scenarios from 2041-2060 (a, c) and 2081-2100 (b, d). The black landmass presents background land that is completely unreachable or unsuitable, while scale of yellow and green landmass represents a scale of less and more suitable regions, respectively.

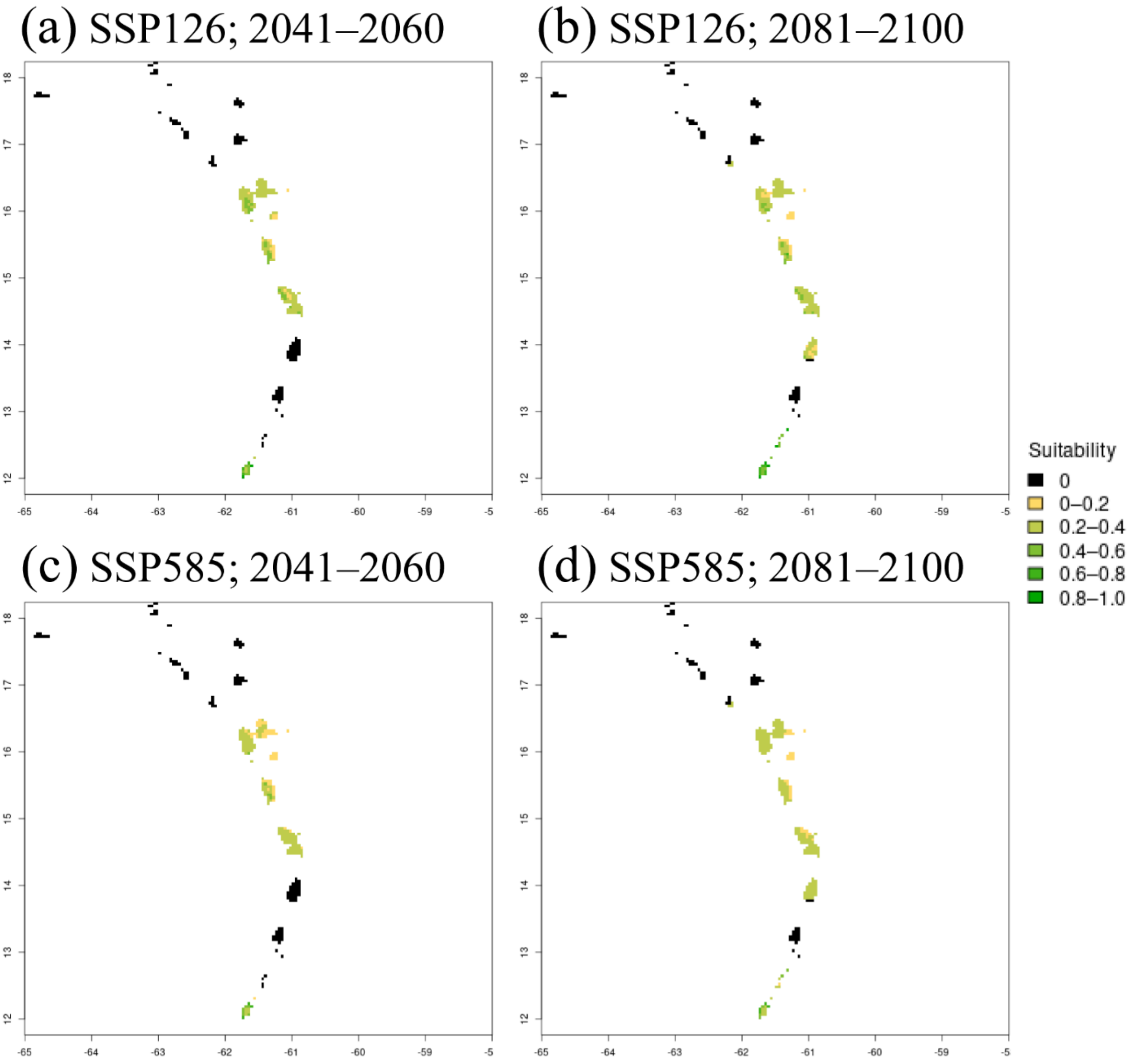


**Fig. S6**: Projected invasive range of *Eleutherodactylus martinicensis* in the Lesser Antilles under low (a, b) and high (c, d) GHG emission scenarios from 2041-2060 (a, c) and 2081-2100 (b, d). The black landmass presents background land that is completely unreachable or unsuitable, while scale of yellow and green landmass represents a scale of less and more suitable regions, respectively.

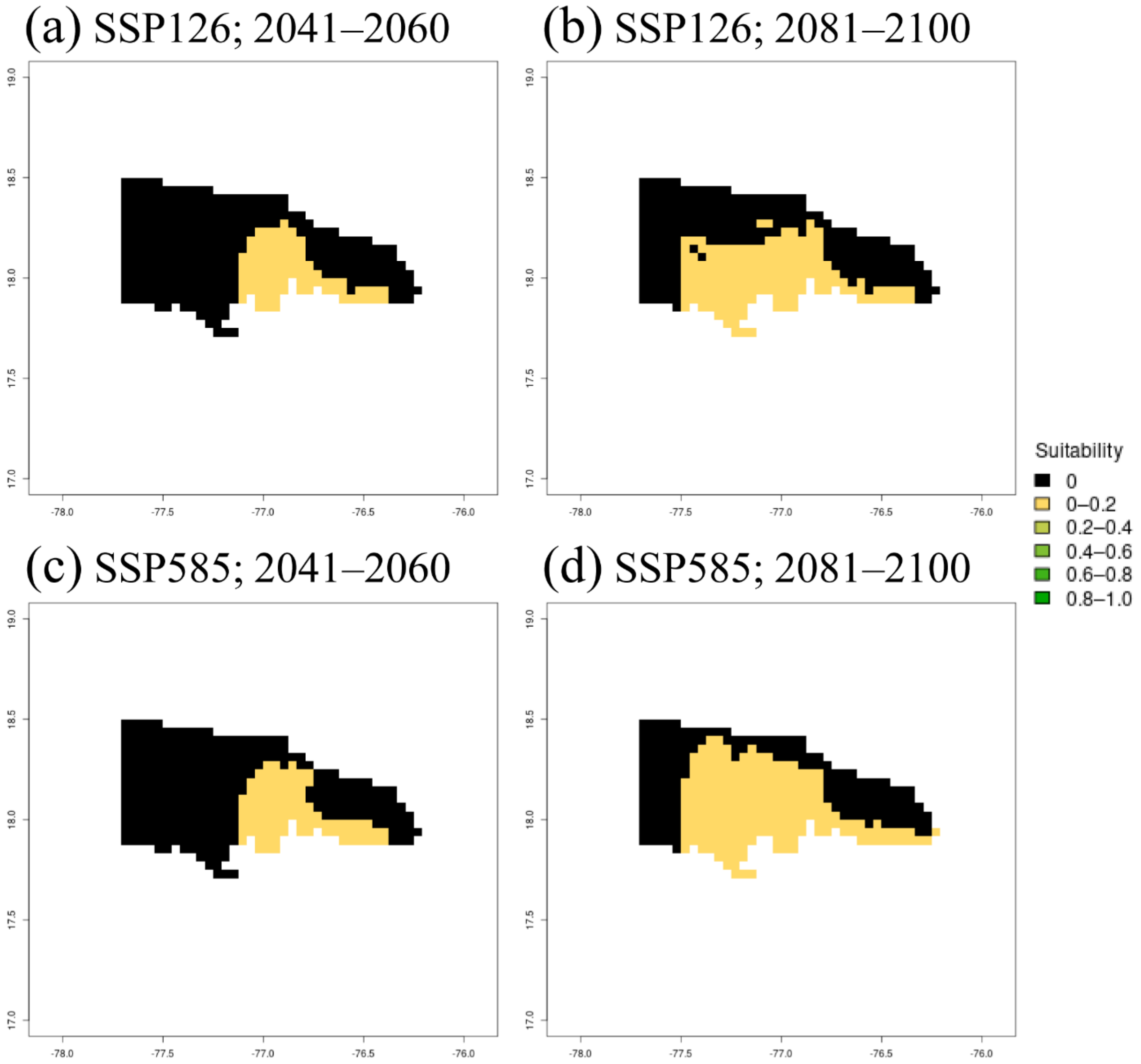


**Fig. S7**: Projected invasive range of *Eleutherodactylus martinicensis* in Jamaica under low (a, b) and high (c, d) GHG emission scenarios from 2041-2060 (a, c) and 2081-2100 (b, d). The black landmass presents background land that is completely unreachable or unsuitable, while scale of yellow and green landmass represents a scale of less and more suitable regions, respectively.

**Table S1:** Maxent ensemble model validation statistics for each species.

| Species | Spatial Rarefying | Feature Class | Regularization Multiplier | BOYCE Model Score | TSS Model Score | ROC Model Score |
|---|---|---|---|---|---|---|
| *Eleutherodactylus coqui* | 559/3,010 | 'linear, quadratic, and hinge' | 1.5 | 0.987 | 0.704 | 0.926 |
| *Eleutherodactylus planirostris* | 2,177/5,019 | 'hinge' | 3.5 | 0.999 | 0.681 | 0.913 |
| *Eleutherodactylus johnstonei* | 218/798 | 'linear, quadratic, and hinge' | 0.5 | 0.955 | 0.572 | 0.879 |
| *Eleutherodactylus antillensis* | 184/1,196 | 'linear' | 0.5 | 0.850 | 0.372 | 0.714 |
| *Eleutherodactylus martinicensis* | 49/302 | 'linear, quadratic, and hinge' | 0.5 | 0.811 | 0.697 | 0.914 |

**Table S2:** Ensemble variable importance for each principal component for each species. Any PC axes denoted as 'NA' fell outside of the 95% window of variance, and thus were not included in the model, for the particular species.

| **Species** | **PC1** | **PC2** | **PC3** | **PC4** | **PC5** | **PC6** | **PC7** | **PC8** |
|---|---|---|---|---|---|---|---|---|
| *Eleutherodactylus coqui* | 0.419 | 0.141 | 0.851 | 0.489 | 0.063 | 0.007 | 0.099 | NA |
| *Eleutherodactylus planirostris* | 0.179 | 0.114 | 0.100 | 0.081 | 0.263 | 0.071 | 0.152 | 0.197 |
| *Eleutherodactylus johnstonei* | 0.876 | 0.043 | 0.418 | 0.150 | 0.612 | 0.567 | NA | NA |
| *Eleutherodactylus antillensis* | 0.232 | 0.430 | 0.575 | 0.026 | 0.005 | 0.312 | NA | NA |
| *Eleutherodactylus martinicensis* | 0.090 | 0.027 | 0.000 | 0.000 | 0.838 | 0.082 | NA | NA |

**Table S3**: Scaled principal component scores for each bioclimatic variable for each species based on native and established datapoints and associated pseudoabsences.

| | ***Eleutherodactylus coqui*** | | ***Eleutherodactylus planirostris*** | | ***Eleutherodactylus johnstonei*** | | ***Eleutherodactylus antillensis*** | | ***Eleutherodactylus martinicensis*** | |
|---|---|---|---|---|---|---|---|---|---|---|
| | **PC1** | **PC2** | **PC1** | **PC2** | **PC1** | **PC2** | **PC1** | **PC2** | **PC1** | **PC2** |
| BIO1 | 0.67 | 0.68 | -0.62 | 0.76 | 0.87 | 0.48 | 0.79 | -0.57 | -0.90 | 0.39 |
| BIO2 | -0.83 | 0.028 | 0.66 | 0.31 | -0.75 | 0.00055 | -0.48 | -0.67 | -0.54 | -0.59 |
| BIO3 | 0.71 | 0.19 | -0.55 | 0.20 | 0.55 | -0.62 | -0.22 | -0.47 | -0.42 | 0.013 |
| BIO4 | -0.89 | -0.15 | 0.82 | -0.13 | -0.69 | 0.31 | -0.55 | -0.0028 | -0.28 | -0.31 |
| BIO5 | -0.58 | 0.51 | 0.065 | 0.86 | 0.70 | 0.65 | 0.41 | -0.86 | -0.95 | 0.16 |
| BIO6 | 0.91 | 0.38 | -0.84 | 0.49 | 0.95 | 0.27 | 0.91 | 0.079 | -0.79 | 0.61 |
| BIO7 | -0.93 | -0.093 | 0.87 | 0.031 | -0.80 | 0.24 | -0.53 | -0.67 | -0.43 | -0.68 |
| BIO8 | 0.15 | 0.71 | -0.57 | 0.41 | 0.78 | 0.57 | 0.81 | -0.50 | -0.87 | 0.35 |
| BIO9 | 0.65 | 0.47 | -0.40 | 0.74 | 0.92 | 0.37 | 0.80 | -0.47 | -0.86 | 0.46 |
| BIO10 | -0.14 | 0.66 | -0.20 | 0.83 | 0.80 | 0.57 | 0.77 | -0.56 | -0.91 | 0.35 |
| BIO11 | 0.86 | 0.48 | -0.79 | 0.59 | 0.92 | 0.37 | 0.83 | -0.51 | -0.89 | 0.42 |
| BIO12 | 0.81 | -0.38 | -0.81 | -0.48 | 0.83 | -0.42 | -0.78 | -0.44 | 0.87 | 0.29 |
| BIO13 | 0.78 | -0.17 | -0.71 | -0.39 | 0.75 | -0.21 | -0.74 | -0.45 | 0.70 | 0.21 |
| BIO14 | 0.49 | -0.49 | -0.55 | -0.43 | 0.59 | -0.64 | -0.47 | -0.015 | 0.70 | 0.43 |
| BIO15 | 0.010 | 0.36 | 0.24 | 0.34 | -0.68 | 0.32 | -0.14 | -0.39 | -0.54 | -0.26 |
| BIO16 | 0.85 | -0.28 | -0.72 | -0.41 | 0.75 | -0.22 | -0.79 | -0.43 | 0.78 | 0.23 |
| BIO17 | 0.61 | -0.50 | -0.59 | -0.47 | 0.63 | -0.63 | -0.62 | -0.15 | 0.80 | 0.40 |
| BIO18 | 0.53 | -0.22 | -0.53 | -0.47 | 0.31 | -0.37 | -0.76 | -0.37 | 0.74 | 0.092 |
| BIO19 | 0.56 | -0.41 | -0.54 | -0.31 | 0.64 | -0.30 | -0.59 | -0.22 | 0.81 | 0.36 |
